# Ciliary Generation of a Peptidergic Sexual Signal

**DOI:** 10.1101/2021.11.08.467802

**Authors:** Raj Luxmi, Richard E. Mains, Betty A. Eipper, Stephen M. King

## Abstract

Peptidergic intercellular communication occurs throughout the eukaryotes, and regulates a wide range of physiological and behavioral responses. Cilia are sensory and secretory organelles that both receive information from the environment and transmit signals. Cilia derived vesicles (ectosomes), formed by outward budding of the ciliary membrane, carry enzymes and other bioactive products; this process represents an ancient mode of regulated secretion. Our previous study revealed the presence of the peptide amidating enzyme, peptidylglycine α-amidating monooxygenase (PAM), in cilia and its key role in ciliogenesis. Furthermore, PAM and its amidated products are released in ciliary ectosomes from the green alga C*hlamydomonas reinhardtii*. One amidated product (GATI-*amide*) serves as a chemotactic modulator for *C. reinhardtii* gametes, attracting *minus* gametes while repelling *plus* gametes. Here we dissect the complex processing pathway that leads to formation of this amidated peptidergic sexual signal specifically on the ectosomes of *plus* gametes. We also identify a potential prohormone convertase that undergoes domain rearrangement during ectosomal secretion as a substrate for PAM. Analysis of this pathway affords insight into how single-celled organisms lacking dense core vesicles engage in regulated secretion, and provides a paradigm for understanding how amidated peptides that transmit sexual and other signals through cilia are generated.

## Introduction

Cilia are membrane-delimited, microtubule-based cell extensions that protrude into the extracellular space and function as key motile, sensory and secretory organelles in many eukaryotes (Marshall and Basto, 2017). These complex organelles that were present in the last eukaryotic common ancestor both receive and transmit signals (Carvalho-Santos et al., 2011; Malicki and Johnson, 2017). Proteins encoded by approximately 5% of the human genome contribute to their assembly, structure and function (van Dam et al., 2019). Mutations in many of these genes cause ciliopathies, with phenotypes ranging from neurological malformations, skeletal abnormalities and kidney disease to obesity and insulin resistance (Reiter and Leroux, 2017). The ciliary localization of receptors for peptides such as Wnt, Hedgehog, insulin, somatostatin and α-melanocyte stimulating hormone (αMSH) plays an essential role in their signaling ability (Anvarian et al., 2019; Green et al., 2016; Wang et al., 2021).

The biosynthesis of signaling peptides involves a well-described sequence of post-translational modifications and proteolytic cleavages that occur as the preproproteins transit from their site of synthesis in the lumen of the endoplasmic reticulum (ER) to the Golgi complex, and are packaged for secretion. Post-translational modifications including disulfide bond formation, N- and O-glycosylation, lipidation, endo- and exo-proteolysis are often required and must occur before secretion (Matsubayashi, 2011; Yasuda et al., 2013). Peptidylglycine α-amidating monooxygenase (PAM), an ancient copper-dependent monooxygenase, catalyzes the final step in the biosynthesis of a broad array of peptides such as mammalian αMSH and vasopressin, the sea urchin sperm attractant resact, as well as numerous invertebrate venom peptide toxins. Amidation occurs *via* a two-step reaction catalyzed by the sequential actions of the monooxygenase (peptidylglycine α-hydroxylating monooxygenase; PHM) and lyase (peptidyl-α-hydroxyglycine α-amidating lyase; PAL) catalytic cores of PAM (Luxmi et al., 2021).

Studies in *Chlamydomonas reinhardtii*, a chlorophyte green alga, revealed the presence of active PAM in the ciliary membrane and demonstrated its key role in ciliogenesis. The ciliary localization of PAM is conserved in mammals, and a role for PAM in ciliary formation and maintenance is observed in mice, zebrafish and planaria (Kumar et al., 2016a; Kumar, 2017; Kumar et al., 2018). Although the catalytic activity of PAM plays an essential role supporting ciliogenesis, how an enzyme essential for a final step in the biosynthesis of secreted peptides contributes to ciliogenesis remains unclear.

*C. reinhardtii* has served as a key model organism for dissecting ciliary assembly, function and signaling (Kumar et al., 2019; Sasso et al., 2018). Although secretory granules that store bioactive peptides in neurons and endocrine cells are not observed, we searched for evidence that *C. reinhardtii* produces and secretes amidated peptides. In addition to constitutive secretion of soluble cargo from Golgi-derived vesicles, the cilia of *C. reinhardtii* shed extracellular vesicles (ectosomes) produced by outward budding from the ciliary membrane (Cao et al., 2015; Long and Huang, 2020; Wang et al., 2014; Wood et al., 2013). Ectosomes released from the cilia of vegetative cells are bioactive and contain a subtilisin-like endoprotease (VLE1) that degrades the mother cell wall (Kubo et al., 2009; Long et al., 2016; Wood et al., 2013). Under plentiful nutrient conditions, haploid *C. reinhardtii* cells divide by mitosis. Starvation triggers a genetically encoded developmental process resulting in the formation of sexual gametes of opposite mating type (termed *minus* and *plus*). Ectosome release increases rapidly when *minus* and *plus* gametes are mixed (Cao et al., 2015). The interaction of their cilia triggers a complex intraciliary signaling pathway that leads to loss of gametic cell walls, formation of mating structures, and cell fusion, yielding a quadriciliate cell that ultimately develops into a diploid zygote. When nutrient conditions improve, the zygote hatches, releasing haploid meiotic progeny (Harris, 2009).

Mass spectrometric analysis of mating ciliary ectosomes led to the identification of an amidated peptide, derived from Cre03.g204500, that acts as a chemotactic modulator, attracting *minus* gametes while repelling *plus* gametes (Luxmi et al., 2019). Amidated peptides from echinoderms, *Hydra*, vespids and humans have also been reported to induce chemotaxis (Palma, 2006; Rowe and Elphick, 2012; Szabó et al., 2015; Takahashi et al., 1997). Cre03.g204500 encodes a 93-kDa protein with all of the features expected of a prepropeptide (hereafter referred to as preproGATI) (Luxmi et al., 2019). Acting on the proprotein (proGATI) created by removal of the signal sequence, a carboxypeptidase B-like enzyme could remove three Arg residues, thus generating a substrate for PAM and production of an amidated C-terminus ending -Gly-Ala-Thr-Ile-NH_2_ (GATI-NH_2_).

The *C. reinhardtii* genome encodes many proteins with the characteristics of prepropeptides. Mating ciliary ectosomes contain proteins derived from several of these prepropeptides, along with the subtilisin-like enzymes needed for their cleavage, PAM and multiple amidated products (Luxmi et al., 2019). C-terminal amidation is often required for peptide bioactivity, as it can greatly enhance affinity for the cognate receptor and confers resistance to proteolytic degradation (Luxmi et al., 2021). Our data suggest that the mating type-specific production and release of bioactive products in ciliary ectosomes represent an evolutionarily ancient path to achieving their regulated secretion.

Here we define the complex processing and amidation pathway leading to formation of the *C. reinhardtii* chemotactic sexual signal and determine how it is trafficked through cilia and ultimately released into ciliary ectosomes and the soluble secretome. We also find that one potential ciliary-localized prohormone convertase is itself a PAM substrate and undergoes an alteration in domain organization during ciliary trafficking, coincident with release of the peptidergic sexual signal. Analysis of proGATI, which yields bioactive products, provides a route to understanding how regulated secretion can occur in a single celled organism lacking peptide storage vesicles. As PAM, peptide processing enzymes, and cilia are broadly conserved in eukaryotes, this study provides a paradigm for understanding how amidated products that transmit chemotactic sexual and other signals through cilia can be generated.

## Results

### Mating ectosomes contain proGATI along with N-terminal and C-terminal fragments of proGATI

Tryptic peptides derived from preproGATI, the protein encoded by Cre03.g204500 and consisting of 908 residues, were identified in both mating ectosomes and the soluble secretome (Luxmi et al., 2018; Luxmi et al., 2019). Interestingly, a proGATI peptide that had been α-amidated and terminated with the sequence -Gly-Ala-Thr-Ile-amide (GATI-*amide*) was identified in mating ectosomes (Luxmi et al., 2019) and in one of six secretome samples analyzed previously (Luxmi et al., 2018). Removal of the N-terminal signal sequence from preproGATI would yield proGATI, with a calculated molecular mass of 90.6 kDa. The amidation of proGATI requires the removal of three Arg residues by a carboxypeptidase B-like exoprotease (Fig. 1A), generating a Gly-extended protein that can serve as a PAM substrate; following α-hydroxylation of this Gly residue by PHM, PAL-mediated cleavage produces a protein terminating with a C-terminal Ile-amide (Fig. 1A).

**Figure 1.**
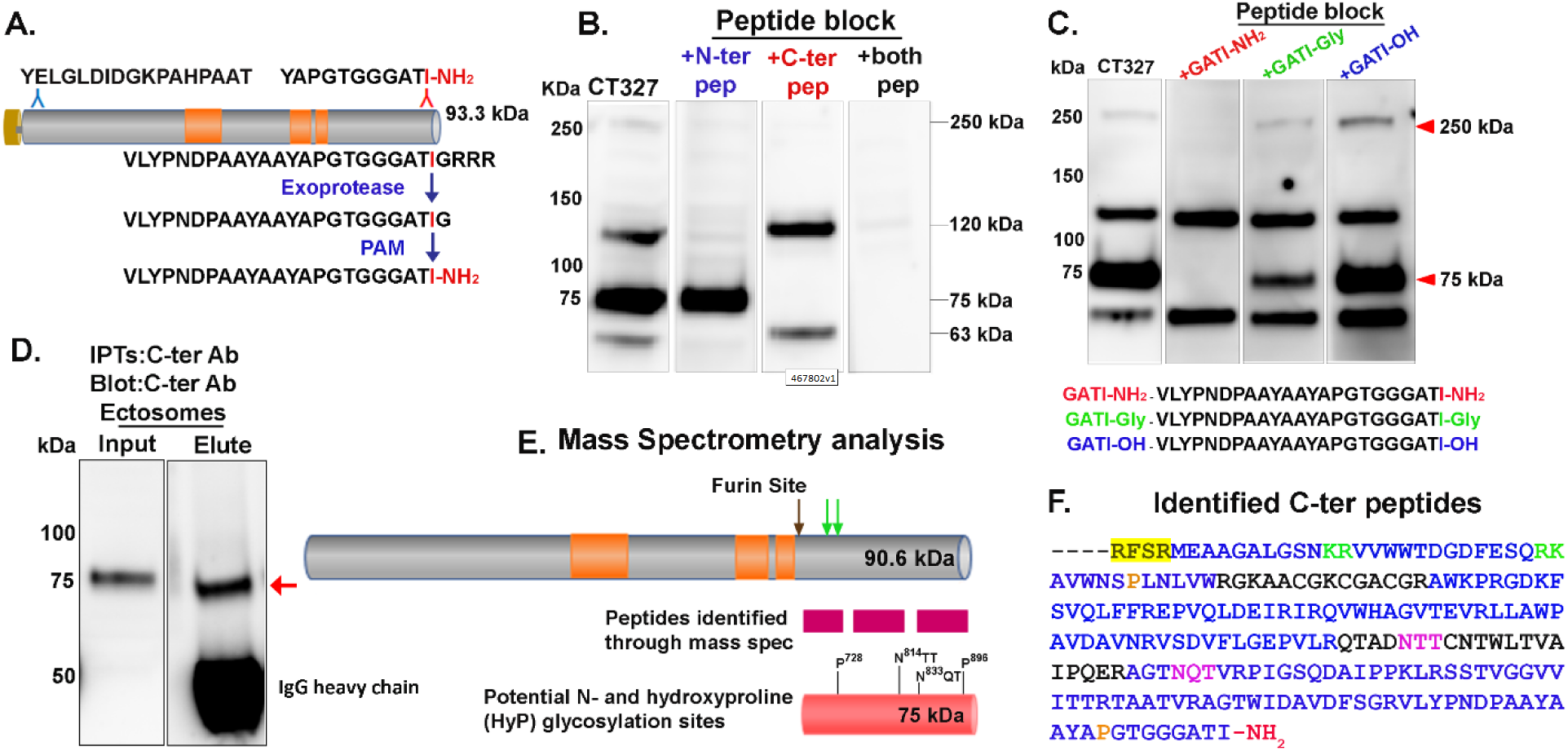
Cre03.g204500 in mating ectosomes. **A**. Diagram shows Cre03.g204500 (preproGATI) and the N-terminal and C-terminal peptides used as antigens and for peptide blocking. The Pro-rich regions are in orange. The pathway leading to C-terminal amidation is illustrated. **B**. Mating ectosomes (10 μg protein) isolated from 1 h mixed gametes were fractionated by SDS-PAGE, blotted and incubated with antiserum (CT327) alone or following pre-incubation with the N-ter (blue), C-ter (red) or mixture of both (black) antigenic peptides. Approximate molecular masses are shown. Data are representative of three independent experiments. **C**. The proGATI antibody generated is amidation specific. Immunoblot of mating ectosomes (10 µg/lane) probed with CT327 antiserum pre-incubated with peptides having -NH_2_ (GATI-amide), –Gly (GATI-Gly) or –OH (GATI-OH) at the C-terminus. Red arrowheads indicate that the signals for the 250-kDa and 75-kDa bands are almost completely blocked by GATI-NH_2_ peptide, attenuated by GATI-Gly and unaffected by GATI-OH. Similar results were obtained in two independent experiments. **D**. Immunoprecipitation from mating ectosomes with affinity-purified C-ter antibody. The excised 75-kDa fragment (red arrow) was analyzed by mass spectrometry. **E**. The location of peptides identified by mass spectrometry is indicated (pink boxes). A furin-like cleavage site (black arrow) precedes the most N-terminally located peptide identified; potential paired basic cleavage sites (green) and predicted N-glycosylation and O-glycosylation sites are indicated. **F**. The C-terminal sequence of proGATI is shown. Peptides identified by mass spectrometry are in blue. The furin-like cleavage site (yellow highlight), paired basic residues (green), predicted N-glycosylation sites (pink), predicted O-glycosylation sites (Pro residues subject to hydroxylation; orange) and amidated C-terminus (red) are indicated.

To explore the biosynthesis, post-translational processing, trafficking and secretion of products generated by the endoproteolytic cleavage of proGATI, we prepared antibodies to a synthetic peptide located near its N-terminus (N-ter peptide) and to a peptide that included the amidated C-terminus (C-ter peptide) (Fig. 1A). Three rabbits were injected with a mixture of carrier-conjugated synthetic peptides and mating ectosomes were used to evaluate the sera. Bands of similar apparent molecular mass were visualized in varying amounts by all three sera; the most prominent appeared at ∼250 kDa, ∼120 kDa, ∼75 kDa and ∼63 kDa (Fig. 1B); post-translational modifications such as N-glycosylation and O-glycosylation can have a dramatic effect on the apparent molecular mass of proteins (Bollig et al., 2007; Voigt et al., 2007).

To test whether these bands were specific, sera were pre-incubated with N-ter peptide, amidated-C-ter peptide or a mixture of both. Pre-incubation with N-ter peptide eliminated the 250-kDa, 120-kDa and 63-kDa signals. Pre-incubation with the amidated-C-ter peptide eliminated the 250-kDa signal and the 75-kDa signal (Fig. 1B). The ability of both peptides to block the appearance of the 250-kDa band suggests that it is an extensively modified form of proGATI. The presence of multiple smaller products indicates that proGATI is subjected to endoproteolytic cleavage.

To determine if the signal produced by the C-ter antibody required amidation, serum was pre-incubated with the amidated C-ter peptide (GATI-NH_2_), the Gly-extended peptide (GATI-Gly) or GATI-OH, which has a free carboxyl group at its C-terminus (Fig. 1C). For both the 250-kDa proGATI band and the 75-kDa band, the signal was greatly reduced by pre-incubation with the GATI-NH_2_ peptide, partially attenuated by pre-incubation with the GATI-Gly peptide and unaffected by the GATI-OH peptide. These data indicate that at least a fraction of the 250-kDa proGATI and 75-kDa product in mating ectosomes is amidated (Fig. 1C).

Affinity-purification was used to prepare antibodies that recognized either the N-terminal region or the amidated C-terminus of proGATI. In agreement with the peptide blocking experiments, affinity-purified N-ter antibody recognized the 250-kDa, 120-kDa and 63-kDa bands in mating ectosomes while C-ter antibody affinity-purified using the GATI-NH_2_ peptide recognized the 250-kDa and 75-kDa bands (Fig. S1A). The specificity of the affinity-purified antibodies was quantified using solid phase assays (Figs. S1B and C).

These data suggest that the 250-kDa protein visualized by both antibodies is a heavily modified version of proGATI, a significant fraction of which is α-amidated. Endoproteolytic cleavage could generate an amidated 75-kDa C-ter fragment along with a 120-kDa N-ter fragment. An additional cleavage could yield a 63-kDa N-ter fragment along with a fragment that would not be recognized by either antibody.

### Endoproteolytic cleavage of proGATI generates a heavily glycosylated 75 kDa product that contains the amidated chemomodulatory peptide

We next used mass spectrometry to identify the proGATI region included in the amidated 75-kDa C-ter fragment immunoprecipitated from mating ectosomes (Fig. 1D). Analysis of in-gel tryptic digests revealed its complete C-terminal amidation. The other tryptic peptides identified provided almost complete (79.7%) coverage of the region from a candidate furin-like cleavage site (R^693^FSR) to Ile^904^-NH_2_, the amidated C-terminus of the chemomodulatory peptide. The calculated polypeptide mass of this cleavage product is 23 kDa (Fig. 1E and F).

Amongst the post-translational modifications that could generate a 23-kDa C-ter proGATI protein backbone with an apparent molecular mass of 75 kDa are N- and O-glycosylation. As in all eukaryotes, N-glycosylation in *C. reinhardtii* involves the assembly of a lipid-linked oligosaccharide that is transferred to target Asn residues in the lumen of the ER followed by maturation in the Golgi complex (Mathieu-Rivet et al., 2020). However, lacking several of the enzymes required for the synthesis of a canonical lipid-linked oligosaccharide, *C. reinhardtii* N-glycans have a unique core structure (Mathieu-Rivet et al., 2020). Two N-glycosylation sites are predicted in this polypeptide using the NetNGlyc tool (Fig. 1E and F). While most of the O-glycans identified in mammals are attached to Ser or Thr residues, in *C. reinhardtii* they are more often attached to hydroxyproline (HyP) residues (Bollig et al., 2007; Joshi et al., 2018; Mathieu-Rivet et al., 2020); two predicted sites (Pro^728^, Pro^896^) occur in the 23 kDa C-ter proGATI region (Figs. 1E and F, and S1D).

Treatment of mating ectosomes with PNGase F, which removes many mammalian N-linked sugars, reduced the apparent molecular mass of a small fraction of the 75-kDa product detected by the C-ter antibody (Fig. S1E); treatment with an O-glycosidase/neuraminidase cocktail was without effect. The mobility of the 120-kDa proGATI fragment recognized by the N-ter antibody was unaltered by either treatment (Fig. S1E). The non-canonical lipid-linked oligosaccharide identified in *C. reinhardtii*, along with its lack of N-acetylglucosaminyltransferase I, which is required for maturation of N-linked oligosaccharides, likely compromise the efficacy of PNGase F; the unique composition of *C. reinhardtii* O-glycans would limit the efficacy of the O-glycosidase cocktail used (Joshi et al., 2018; Mathieu-Rivet et al., 2020).

### HEK-293 cells synthesize and secrete heavily glycosylated, amidated proGATI

To facilitate our understanding of the proGATI protein and the modifications involved in producing the amidated 75 kDa protein secreted in mating ectosomes, we stably expressed a cDNA encoding preproGATI in a human embryonic kidney cell line (HEK-293). In *C. reinhardtii*, as in other species, maturation of newly synthesized glycoproteins and their ability to move from the ER into the Golgi are monitored by their interactions with calnexin and calreticulin (Mathieu-Rivet et al., 2020). We reasoned that the efficient secretion of proGATI by HEK-293 cells would indicate proper folding and allow usage of tools available to study vertebrate N- and O-glycosylation. Affinity-purified C-ter proGATI antibodies were used to evaluate cell extracts and spent medium. Specific bands of 120 kDa and 170 kDa were detected in cell extracts by both N-ter and C-ter antibodies (Figs. 2A and S2A). A minor doublet at ∼37-kDa was also detected in cell extracts, but not in spent medium (Fig. 2A). The fact that spent medium contains a single 170-kDa protein recognized by both N-ter and C-ter antibodies led to its identification as HEK-proGATI; differences in the N- and O-linked oligosaccharides attached to proGATI produced by HEK cells and by *C. reinhardtii* would account for the difference in apparent molecular mass.

**Figure 2.**
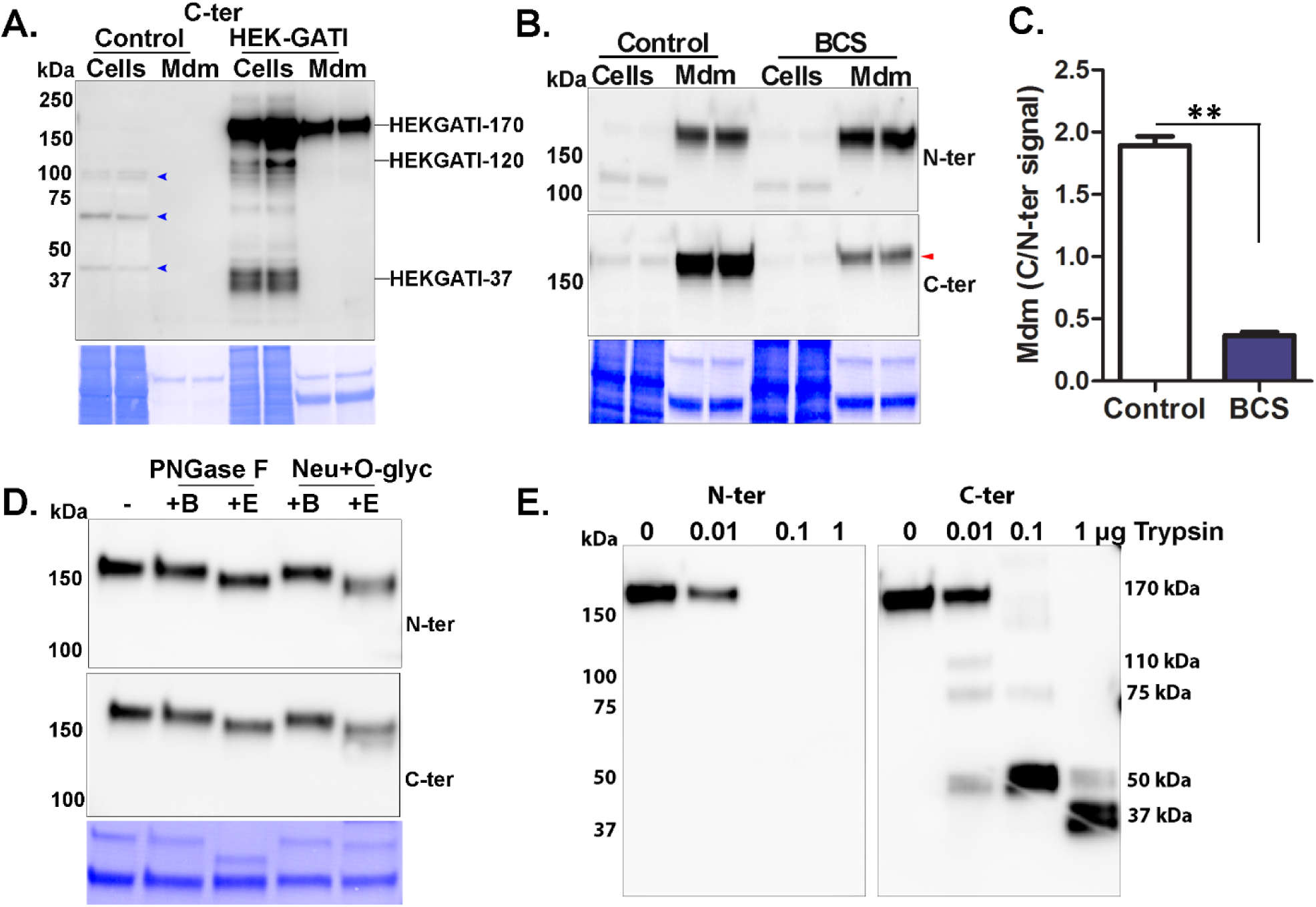
Expression of proGATI in HEK-293 cells. **A**. Immunoblot of cell extracts (Cells; 20 μg protein, approximately 10 % of total) and spent medium (Mdm; 1% of total collected over an 18 h period) of Control (non-transfected) and HEK-293 cells expressing preproGATI probed with affinity-purified C-ter antibody. A 170 kDa (HEK-proGATI) band was detected in both cells and spent medium while 120 kDa and 37 kDa bands were detected only in cells (and see Fig. S2A). Non-specific bands identified in Controls are marked (blue arrow). Secretion rate and cell content are quantified in Fig. S2B. **B**. Analysis of C-terminal amidation of HEK-proGATI. Spent medium (5%) and cell lysates (15 µg, ∼20% of total) of BCS-treated cells and their respective Controls were analyzed. The C-ter signal for HEK-proGATI was reduced following BCS treatment (red arrow), whereas the N-ter signal was unaffected. **C**. The C-ter/N-ter signal ratio for HEK-GATI-170 was reduced following BCS treatment. Results are the average of duplicates, where **P<0.001. **D**. HEK-proGATI spent medium (10 µl) was digested with PNGase F or with a mixture of O-glycosidase and neuraminidase (Neu+O-glyc); no treatment (-), incubated in buffer alone (+B), or buffer with enzyme (+E). Reductions in the apparent molecular mass of secreted HEK-GATI-170 of ∼15-20 kDa were observed. Similar results were obtained in three independent experiments. **E**. Tryptic digestion of HEK-GATI in spent medium (10 µl); samples were fractionated by SDS-PAGE and probed with N-ter and C-ter antibodies. The N-ter antigenic site contains a single Lys residue and is destroyed by trypsin treatment. The C-ter antibody detected the indicated tryptic products. The results were duplicated in independent experiments.

To test whether HEK-293 cells amidate proGATI, bathocuproine disulfonate (BCS) was used to deplete cellular copper, inhibiting the activity of the amidating enzyme, PAM (Bonnemaison et al., 2015). While the 170-kDa N-ter signal was unaltered following BCS treatment, the 170-kDa C-ter signal fell dramatically (Fig. 2B). To account for any differences in secretion rate, the ratio of 170-kDa C-ter signal to 170-kDa N-ter signal was calculated. BCS treatment caused a four-fold reduction in this ratio, consistent with the conclusion that HEK-293 cells amidate the C-terminus of the proGATI that they secrete (Fig. 2C).

ProGATI includes six potential N-glycosylation sites (Asn-X-Ser/Thr) and several potential O-glycosylation sites (-Ser/Thr and HyP) (Fig. S1D). Digestion with either PNGase F or a mixture of O-glycosidase and neuraminidase reduced the apparent molecular mass of secreted HEK-proGATI by ∼15-20 kDa, consistent with the occurrence of extensive N- and O-glycosylation (Fig. 2D).

Successful ectosome-mediated delivery of a chemomodulatory peptide such as GATI-NH_2_ would require it to be resistant to proteolysis. Spent medium containing HEK-proGATI was used to test this hypothesis. Exposure to increasing amounts of trypsin eliminated the N-ter signal and generated a series of smaller products recognized by the C-ter antibody (Fig. 2E). Cleavage at the single Lys residue in the N-ter peptide is consistent with this result (Fig. 1A). Trypsin produced a sequence of smaller products detected by the C-ter antibody. C-ter signal intensity was not diminished, with essentially complete conversion of 170 kDa HEK-proGATI into a 50 kDa and then a 37-kDa product, which may resemble the amidated 75 kDa C-ter fragment found in mating ectosomes (Fig. 2E).

### Purification and domain organization of proGATI

Since HEK-proGATI is amidated and secreted rapidly (Fig. S2B), we undertook its purification from spent medium (Fig. 3A and S2C) and analysis using mass spectrometry. Although the N- and O-glycans attached to HEK-proGATI will differ from those attached to proGATI produced by *C. reinhardtii*, the sites available to enzymes involved in N- and O-glycosylation are expected to be the same. Purified native and deglycosylated HEK-proGATI were analyzed, using a cocktail of enzymes designed to remove both N- and O-linked glycans. Deglycosylation reduced its apparent molecular mass by ∼30 kDa (Fig. 3B).

**Figure 3.**
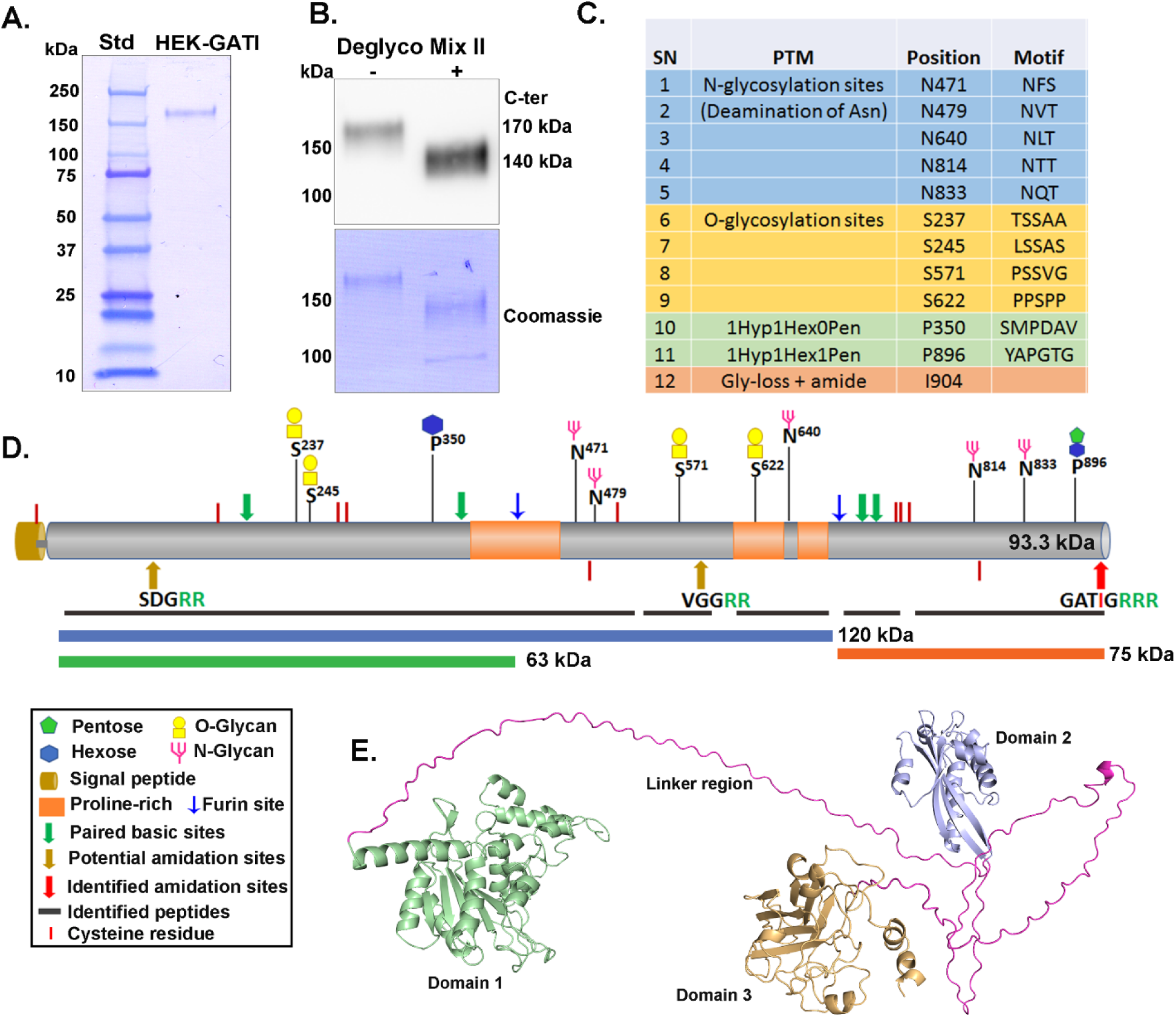
Mass spectrometry analysis of purified HEK-GATI-170. **A**. SDS-PAGE of purified HEK-proGATI (see Fig. S2C); Coomassie-stained PVDF membrane is shown. **B**. Digestion of purified HEK-proGATI with protein deglycosylation mix II reduced its apparent molecular mass. **C**. Glycosylation sites identified; N-glycosylation sites were identified in deglycosylated-HEK-proGATI, while O-glycosylation sites were identified in purified native protein. **D**. Schematic diagram of preproGATI illustrating the N- and O-glycosylation sites identified in purified HEK-proGATI. The predicted products resulting from cleavage at the furin-like sites are indicated. **E**. The structural model of proGATI generated using RoseTTAFold (Baek et al., 2021) contains three well-folded domains (domain 1, residues 51-370, green; domain 2, residues 446-593, blue; domain 3, residues 696-908, orange) connected by long, highly flexible, Pro-rich linkers (pink). Although individual domains are well structured, their relative orientation with respect to each other is variable.

Mass spectrometry of native HEK-proGATI identified four O-glycosylated Ser residues and two O-glycosylated HyP residues. Analysis of enzymatically deglycosylated HEK-proGATI identified five N-glycosylation sites. The amidated C-terminus (-GATI-NH_2_) was found in both samples; C-terminal peptides ending in –Gly and –Gly-Arg were also identified indicating that carboxypeptidase processing and amidation had not gone to completion (Fig. 3C and D). Peptides spanning the entire sequence of HEK-proGATI were identified (87.3% coverage) (Fig. 3D).

The ability of trypsin to convert amidated HEK-proGATI into stable, amidated products as small as 37 kDa (Fig. 2E), is consistent with the presence of stable domains. To explore this possibility, a structural model of proGATI was generated using RoseTTAFold (Baek et al., 2021). The proGATI prediction includes three well-folded domains connected by highly extended, Pro-rich flexible linkers (Fig. 3E). The signal sequence was not included in the structural model. N-terminal domain 1 contains 323 residues (green), terminating just before a Pro-rich region. Domain 2 includes 153 residues and domain 3 has 213 residues, ending at the C-terminus. The 70-residue linker between domains 1 and 2 contains 37 Pro residues and a furin-like site (K^407^PRK), while 43 of the 99 residues in the second linker are Pro residues; these very Pro-rich regions likely contribute to the abnormal migration of proGATI during SDS-PAGE. Domain 3 forms an antiparallel β-sandwich and has a nominal molecular mass of 23 kDa with a pI of 10 (Fig. 3E). This domain corresponds precisely to the C-terminal region identified by mass spectrometry of the 75-kDa amidated product in mating ectosomes, and is immediately preceded by a furin-like cleavage site. Cleavage at this site alone would release domains 1 and 2 (predicted to represent the 120-kDa N-terminal fragment), while further proteolysis at K^407^ might generate the 63-kDa N-terminal product. Domain 3, which includes four Cys residues, has a single predicted disulfide bond (C^739^ and C^745^); although C^742^ and C^817^ are located close to each other, a significant rearrangement would be needed for disulfide bond formation. Importantly, the experimentally confirmed C-terminal amidation site, the Pro residue that is O-glycosylated (P^896^) and both Asn residues that are N-glycosylated (Asn^814^ThrThr and Asn^833^GlnThr) are completely exposed and accessible for modification in the model structure (Fig. S3).

### Ciliary localization and mating type-specific processing of proGATI

Under nutrient deprivation conditions, *C. reinhardtii* cells differentiate into *minus* and *plus* gametes that expresses mating type-specific genes, enabling them to recognize each other. Our previous study showed that CrPAM expression increased during gametogenesis, and that the C-ter antigenic peptide and a longer synthetic amidated peptide (VLYPNDPAAYAAYAPGTGGGATI-NH_2_) produced a mating type-specific chemotactic response, attracting *minus* gametes and repelling *plus* gametes (Luxmi et al., 2019). These observations prompted investigation of proGATI in gametes. Cells, deciliated cell bodies and cilia were subjected to immunoblot analysis. Use of the N-ter and C-ter antibodies revealed enrichment of 250-kDa proGATI in the cilia of both *minus* and *plus* gametes (Fig. 4A). In contrast, the C-ter antibody detected a 75-kDa band only in *plus* gametes; while detectable in *plus* gamete cells, the 75-kDa band was enriched in *plus* gamete cilia and essentially undetectable in deciliated cell bodies. Strikingly, production of amidated 75-kDa GATI is specific to *plus* gametes (Fig. 4A and 4B). Mass spectrometric analysis of 75-kDa GATI immunoprecipitated from *plus* gamete cell lysates confirmed complete amidation of its C-terminus and the presence of peptides like those identified in 75-kDa GATI immunoprecipitated from mating ectosomes (Figs. 1E, F and S4).

**Figure 4.**
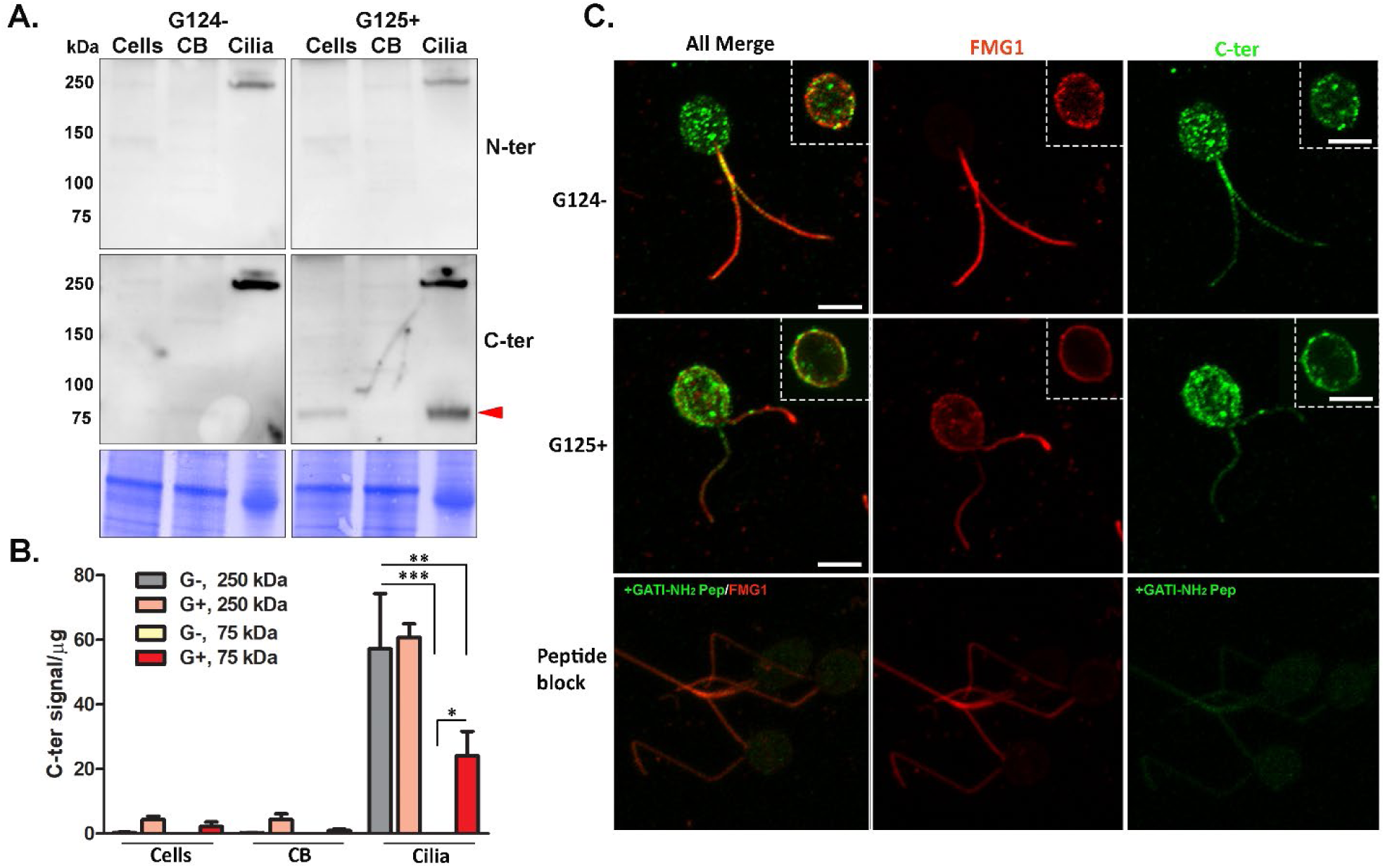
Processing and localization of proGATI in *minus* and *plus* gametes. **A**. Immunoblot of cells, deciliated cell bodies and cilia of *minus* (G124-) and *plus* (G125+) resting gametes using affinity-purified proGATI N-ter and C-ter antibodies. Equal amounts of protein (20 µg) were loaded. **B**. Quantification of the C-ter signal for proGATI revealed significant enrichment of 250-kDa and 75-kDa bands in cilia but not in cell bodies (CB). Results are the average of duplicates. Means were compared with ± range. Asterisks indicate significant differences between groups *P<0.05, **P<0.01, ***P<0.001. **C**. Maximal projection confocal images of *minus* and *plus* resting gametes stained with the C-ter proGATI antibody (green) and antibody to FMG1 (red). Inset images show single Z-planes. *Plus* gametes probed with antibody pre-incubated with the GATI-NH_2_ peptide exhibit reduced staining (green) in cell bodies and cilia. Similar localization of proGATI in gametes was obtained in three independent experiments. Scale bar = 5 µm.

We next used immunofluorescence microscopy to determine the subcellular localization of GATI-derived proteins in resting gametes. Maximal Z-projection confocal images of *minus* and *plus* gametes showed that the C-ter GATI signal was localized in discrete puncta throughout the cytoplasm (Fig. 4C); this signal could represent intact 250-kDa proGATI and/or the 75-kDa C-ter product derived from it. In contrast, simultaneous visualization of FMG1, a ciliary membrane glycoprotein, revealed more signal in the cilia and around the margins of the cell body. Single Z-stack images showed the accumulation of C-ter GATI signal at the cell surface, co-localized with FMG1 (Fig. 4C, inset). Diffuse C-ter GATI signal along the length of the cilia was also observed in both mating types (Fig. 4C). To confirm staining specificity, the C-ter antibody was pre-incubated with antigenic peptide; signal intensity (green) was greatly reduced in the cell body, on the cell surface and in cilia of *plus* gametes (Fig. 4C, lower panel). The punctate staining in the cell body could represent Golgi-derived vesicles, which may enter the ciliary membrane after accumulating on the cell surface.

### Mating triggers ectosomal trafficking and processing of the GATI-precursor

Ectosome formation involves outward budding of the ciliary membrane (Wood et al., 2013). The initiation of mating triggers the formation and release of ectosomes (Cao et al., 2015); the catalytic domains of CrPAM are exposed on the outer surface of mating ectosomes and PAM is not found in the soluble secretome (Kumar et al., 2016a; Luxmi et al., 2019). ProGATI, which lacks a transmembrane domain, is in both mating ectosomes and the secretome. We utilized affinity-purified proGATI antibodies and immunogold-electron microscopy to determine its ectosomal localization.

Ectosomes from mating gametes were embedded in agarose and imaged by thin section transmission EM (Fig. 5A); the vesicles range from ∼80 nm to ∼260 nm in diameter. Following incubation with intact ectosomes, affinity-purified proGATI antibodies were visualized using a gold-tagged anti-rabbit secondary antibody and negative stain EM; signals obtained with both antibodies were localized on the ectosome surface (Fig. 5A). Ectosomes incubated only with gold-conjugated secondary antibody served as a negative control.

**Figure 5.**
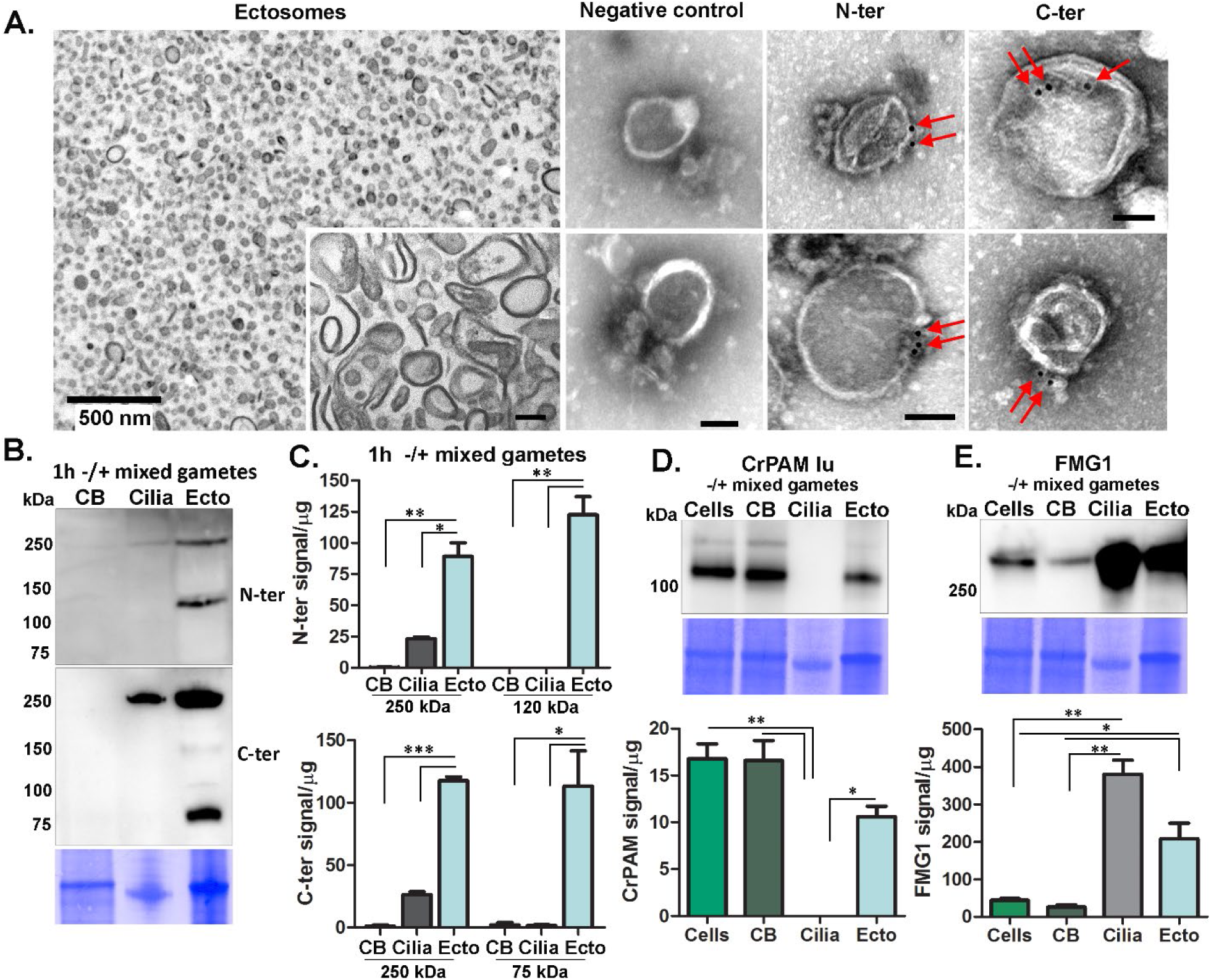
ProGATI in mating ectosomes. **A**. A cross-section transmission electron micrograph of an agarose embedded ectosome pellet isolated from 1 h -/+ mixed gametes is shown. The inset shows a higher magnification image of ectosomes that had been treated with Na_2_CO_3_ to remove peripheral membrane proteins; scale bar = 100 nm. The right panels show immuno-gold-EM negative stain images of intact ectosomes incubated with affinity-purified N-ter or C-ter antibodies and a gold-tagged secondary antibody; both epitopes localized to the ectosomal surface. Ectosomes incubated with gold-tagged secondary anti-rabbit antibody alone served as a negative control; scale bars = 500 nm (main image) and 100 nm (inset). Images are representative of three independent experiments. **B**. The deciliated cell bodies (CB), cilia and ectosomes (Ecto) isolated from mixed gametes were fractionated by SDS-PAGE, blotted and probed with the N-ter and C-ter antibodies against proGATI. **C**. Graph showing the enrichment of N-ter and C-ter signals for proGATI and its fragments in ectosomes. Results are the average of two independent experiments; mean is ± range. Asterisks indicate a statistically significant difference between two groups (*P < 0.05, **P<0.001, ***P<0.0001). **D & E**. Immunoblot analysis showing CrPAM and FMG1 levels in cells, cell bodies, cilia and ectosomes isolated from mixed gametes. Quantification of CrPAM and FMG1 protein levels is shown in the graphs. Results are average of duplicates and error bars indicate the ± range (where *P < 0.05, **P<0.001).

Ectosomes, deciliated mixed gamete cell bodies and cilia prepared 1 h after the initiation of mating were subject to immunoblot analysis (Fig. 5B). Based on use of both the N-ter and C-ter antibodies, cilia contained only 250-kDa proGATI. In contrast, mating ectosomes contained 250-kDa proGATI along with the 120 kDa N-ter fragment and the 75 kDa C-ter fragment, whereas proGATI products were not detected in the cell bodies (Fig. 5B). Quantification revealed enrichment of 250-kDa proGATI in mating ectosomes and an even greater enrichment of both the 120 kDa and 75 kDa fragments (Fig. 5C), suggesting that the cleavage creating them occurs on the ectosomal surface.

For comparison, we evaluated the specificity and selectivity with which two other ciliary proteins, CrPAM and FMG1, move from cilia into ectosomes during mating (Fig. 5D and E). CrPAM was previously identified in mating ectosomes, but was not found in vegetative ectosomes (Luxmi et al., 2019), while FMG1 is present in both (Long et al., 2016; Luxmi et al., 2019). Immunoblot analysis confirmed the presence of both CrPAM and FMG1 in mating ectosomes. After an hour of mating, very little CrPAM remained in the cilia; although mating ectosomes contained CrPAM, its ectosomal levels did not exceed those in the cell body (Fig. 5D). FMG1 levels in cilia and mating ectosomes greatly exceeded those in cell bodies, but FMG1 levels in mating ectosomes did not exceed those in cilia (Fig. 5E). Thus, the cell bodies of mating gametes were essentially devoid of proGATI while both CrPAM and FMG1 were readily detected.

### Differential release of proGATI products from mating ectosomes

We previously found that both CrPAM protein and enzyme activity associate with the ciliary axoneme; this interaction is disrupted by treatment with 0.6 M NaCl following detergent extraction (Kumar et al., 2016a). To explore the ciliary distribution of proGATI and its products, we isolated cilia from resting gametes of both mating types and from 1 h mixed gametes. Isolated cilia were first treated with Triton X-100 to release membrane proteins and soluble matrix components. This was followed by treatment with 0.6 M NaCl to extract proteins that were tightly bound to the axoneme; the resulting extracted axoneme pellet was solubilized in 1× SDS buffer.

The amidated 75-kDa GATI product was detected in the cilia of *plus* but not *minus* gametes (Figs. 6A and B). This fragment was largely recovered in the detergent soluble fraction, with a smaller amount released by 0.6 M NaCl; it was not present in the axonemal fraction of *plus* gametes. The 250-kDa proGATI protein was in the detergent soluble, 0.6 M NaCl and axonemal fractions from both gametes (Figs. 6A and B). During gamete mating, the amidated 75-kDa GATI product and the 250-kDa proGATI protein were released into ectosomes (Figs. 6A and B). The presence of an N-terminal fragment of proGATI in ectosomes but not in the ciliary fractions suggested that cleavage of 250 kDa proGATI occurs on the ectosomal surface.

**Figure 6.**
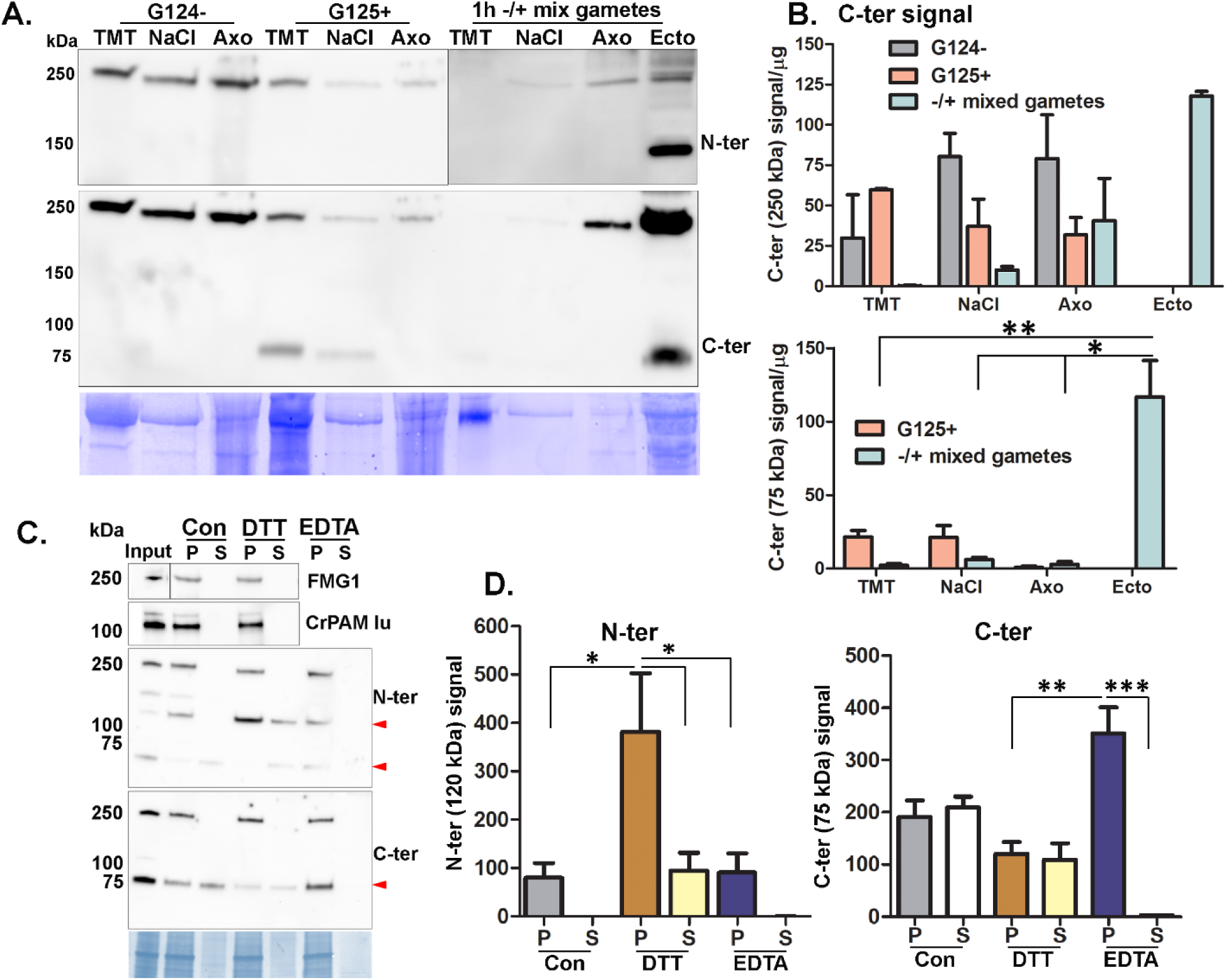
Ciliary localization and association of proGATI and its fragments with ectosomes. **A**. Cilia were sequentially treated with buffers containing 1% Triton X-100 (TMT) and 0.6 M NaCl (NaCl); the resulting axonemal pellet (Axo) was solubilized in 1% SDS-buffer. The sub-ciliary fractions from resting *minus* (G124-) and *plus* (G125+) gametes mixed gametes and mating ectosomes (Ecto) were fractionated by SDS-PAGE, blotted and probed with affinity-purified N-ter and C-ter antibodies. Equal amounts of protein (20 µg) were loaded for each sample. **B**. Immunoblot quantification of the 250-kDa and 75-kDa C-ter products. Means are average of duplicates and error bars indicate ± range, where *P<0.05, **P<0.01. **C**. Freshly isolated mating ectosomes (Input) were washed with buffer alone (10 mM HEPES, control) or with buffer containing 10 mM dithiothreitol (DTT) or 10 mM EDTA; after centrifugation, the resulting supernatants (S) and pellets (P) were analyzed for the presence of proGATI (using N-ter and C-ter antibodies), PAM and FMG1. Red arrowheads mark the 120-kDa, 75-kDa and 63-kDa bands. Samples loaded represent the pellets and corresponding supernatants derived from an initial 15 µg of ectosomes. **D**. Quantification of 120-kDa N-ter signal (n = 4) and 75-kDa C-ter signal (n = 3); means ± SEM are shown. Asterisks indicate significant differences between the groups, *P<0.05, **P<0.01, ***P<0.001.

To evaluate how 250 kDa proGATI (which lacks a transmembrane domain) associates with the ectosome surface, freshly isolated ectosomes were washed with 10 mM HEPES buffer (control) or with buffer containing 10 mM dithiothreitol (DTT) or 10 mM EDTA, and the resulting supernatants examined for the release of GATI products, CrPAM and FMG1 (Figs. 6C and D). Neither 250-kDa proGATI, CrPAM nor FMG1 was solubilized, with signal detected only in the ectosomal pellets. In contrast, the amidated 75-kDa C-terminal product and N-terminal 63-kDa segment were both released by washing with low ionic strength HEPES buffer; release did not occur following chelation of divalent cations with EDTA. Although not released by buffer alone or by EDTA treatment, the N-terminal 120-kDa GATI fragment was partially displaced from ectosomes by 10 mM DTT. This effect was DTT-specific; treatment with 10 mM β-mercaptoethanol had no effect (Fig. S5). These results suggest that all three domains individually mediate associations with the ectosomal surface. This tripartite attachment mechanism likely explains why release of 250-kDa amidated proGATI was not observed under any conditions.

### Distribution, processing and amidation of putative prohormone convertases in cilia

The appearance and accumulation of the 75-kDa amidated proGATI product on *plus* (but not *minus*) gamete cilia, and of both 120- and 75-kDa proGATI products on mating ectosomes, suggested that proteolytic processing occurs on the ciliary and/or ectosomal surface or during the sorting and transit of the precursor from cilia into nascent ectosomes. Mating ectosomes contain two subtilisin-like proteases, SUB14 and VLE1; they are the closest *Chlamydomonas* homologs of mammalian prohormone convertases PC2 and PCSK7, respectively (Luxmi et al., 2019). To address the ciliary distribution of these putative prohormone convertases, we performed comparative proteomics of cilia from vegetative and gametic cells of both mating types. This confirmed the presence of VLE1 in vegetative cilia of both mating types (Kubo et al., 2009); SUB14 was not detectable in vegetative cilia (Fig.7A and Supplemental Data File 1). Strikingly, VLE1 was identified in the cilia of *plus* gametes, but was not detected in *minus* gamete cilia. SUB14 expression was also mating type specific, but it was present in the cilia of *minus*, but not *plus*, gametes (Fig.7A). In consequence, VLE1 is the only putative prohormone convertase present in ciliary samples that contain proteolytically processed proGATI products.

Peptides from the cytosolic, pro-, S8 and C-terminal domains of VLE1 (Fig. 7B) were identified in cilia from vegetative and *plus* gamete cells. In contrast, mating ectosomes and the secretome contained only peptides from the S8 and C-terminal domains (Fig. 7B). Activation of subtilisin-like prohormone convertases generally requires autoproteolytic cleavage and subsequent dissociation of the pro-domain (Shakya and Lindberg, 2020). Clustal analysis identified the –Gly-Arg-Arg site that immediately precedes the catalytic domain as the likely site for autoactivation. Autoproteolytic cleavage at this site, followed by exoproteolytic removal of the two Arg residues would produce an amidation site. Mass spectrometry revealed that all of the ciliary VLE1 had been proteolytically processed at this site and was amidated (Fig.7B); partially processed peptides derived from this region of VLE1 and ending in –Gly, –Gly-Arg or – Gly-Arg-Arg were not observed. Detailed analysis of the predicted VLE1 structure (Fig. 7C) and peptides identified in cilia suggests that the pro-domain remains associated with the S8 domain, tethering it to the ciliary membrane even after autoproteolytic cleavage and amidation.

**Figure 7.**
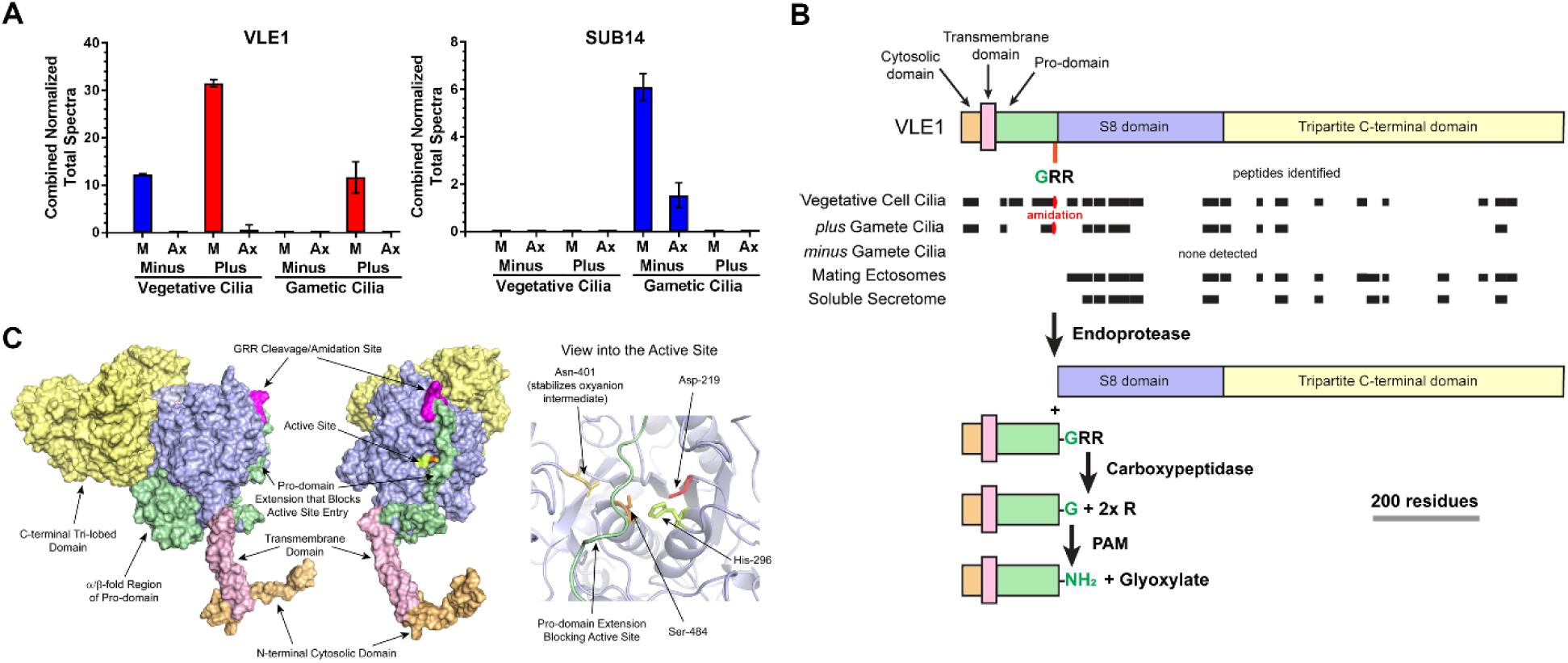
Ciliary distribution and processing of subtilisin-like proteases. **A**. Normalized total spectral counts of VLE1 and SUB14 in the ciliary membrane/matrix (M) and axonemal (Ax) fractions from *minus* and *plus* vegetative and gametic cilia (mean ± SEM; n=3). **B**. Diagram showing the cytosolic (light orange), transmembrane (pink), pro-(green), S8 (purple) and C-terminal (yellow) domains of VLE1. The l VLE1 peptides identified in the cilia of vegetative cells, *plus* and *minus* gametes and in mating ectosome and the soluble secretome are indicated by black boxes. The cleavage/amidation site (GRR) that immediately precedes the S8 catalytic domain is indicated. The processing pathway proposed for VLE1 is shown. The α-amidated peptide (K)APTDITDPTAASSSS-NH_2_ produced by cleavage and amidation was found in both vegetative and *plus* gamete cilia. **C**. Two views of the molecular surface of a structural model for VLE1 calculated using RoseTTAFold are shown. The protein consists of a short N-terminal cytosolic domain (light orange), a single transmembrane region (pink), an unusual pro-domain (green), the catalytic S8 domain (blue), and a large C-terminal domain (yellow) that has a tri-partite organization with each lobe consisting of two anti-parallel β sheets which exhibit considerable structural similarity to the CEA1 N-acetylglucosamine-binding adhesin from the methylotrophic yeast *Komagataella pastoris* (z score = 11.8, RMSD = 4.0 Å; 5A3L). The cleavage/amidation site is indicated in magenta. The right-hand panel shows a ribbon diagram of the active site. Side chains of the catalytic triad residues and the Asn that stabilizes the transition state are shown. The pro-domain strand that arches across the active site is indicated in green.

## Discussion

Identification of an amidated peptide that has a mating-type specific effect on *C. reinhardtii* mobility led us to explore the properties of its putative precursor, the manner in which this precursor might be converted into smaller products, and the regulated secretion of its product peptides.

### ProGATI undergoes extensive post-translational modification and contains multiple domains

The *C. reinhardtii* genome encodes hundreds of proteins with the general characteristics of prepropeptides (Luxmi et al., 2019). As observed in the ER of metazoans, preproGATI undergoes signal peptide removal, along with the first steps of N- and O-glycosylation (Fig. 8A). ProGATI, like many other putative *C. reinhardtii* propeptides, is quite large, with a predicted molecular mass of 90.6 kDa, and multiple domains connected by Pro-rich linker regions. In addition to the Asn and Ser/Thr sites subject to N- and O-glycosylation in metazoan propeptides, hydroxy-Pro residues in domains 1 and 3 of proGATI are O-glycosylated (Fig. 3). In plants and algae, hydroxy-Pro residues are major O-glycosylation sites for addition of pentose (arabinogalactan) sugars (Bollig et al., 2007; Tan et al., 2003). With a unique core structure to their N-glycans and unique O-glycosyl transferases, propeptides synthesized by *C. reinhardtii* differ in important ways from metazoan propeptides (Joshi et al., 2018; Mathieu-Rivet et al., 2020; Schulze et al., 2017; Xu et al., 2020).

**Figure 8.**
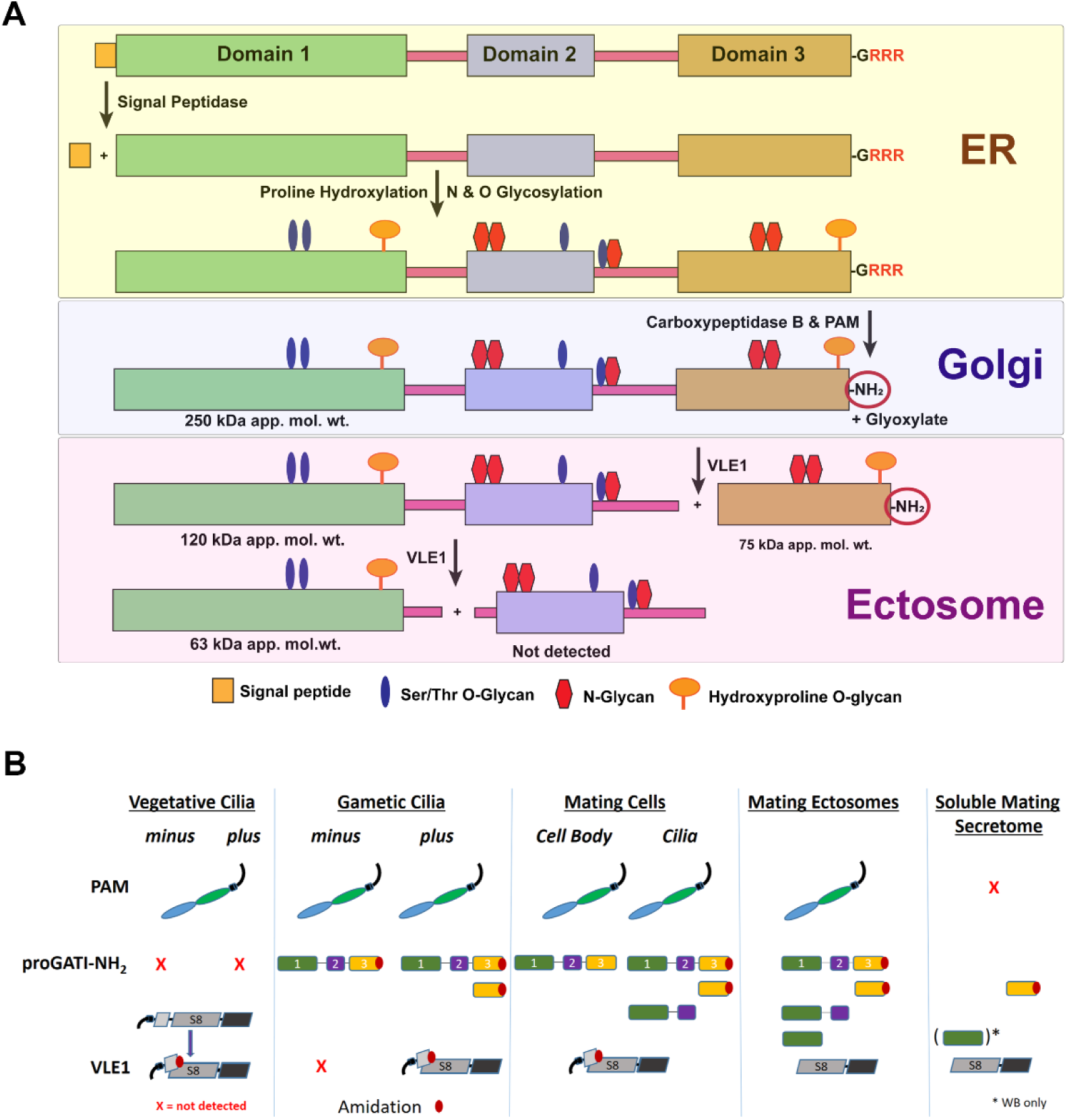
ProGATI processing pathway. **A**. Diagram illustrating the processing pathway of preproGATI that occurs as it traffics through the ER and Golgi and subsequently enters cilia and ectosomes. As preproGATI enters the ER, its signal peptide (orange box) is removed by signal peptidase. The addition of N-linked sugars (red) begins in the ER, as does modification of Pro to HyP by prolyl hydroxylases. As proGATI moves into the Golgi complex, more complex sugars and O-linked sugars on HyP (orange) and Ser/Thr (blue) residues are added, leading to the higher apparent molecular mass (∼250 kDa) of proGATI. A carboxypeptidase trims the three C-terminal Arg residues and generates a substrate for PAM. PAM converts the –Gly extended substrate into the amidated product (GATI-NH_2_) in a two-step reaction and releases glyoxylate as a byproduct. This 250-kDa amidated proGATI form is then moved to the ciliary membrane. Once on cilia, or as it moves from cilia into nascent ectosomes, 250-kDa proGATI is cleaved by a subtilisin-like endoprotease (predicted to be VLE1) to yield the 120-kDa N-terminal region, and the amidated 75-kDa C-terminal fragment. Cleavage of the 120-kDa product at either a second furin-like cleavage site located in the linker between domains 1 and 2, or at a dibasic site at the C-terminal end of domain 1, might then produce the 63-kDa N-terminal fragment and a second product containing domain 2 for which no probe currently exists. **B**. Diagram illustrating the presence and absence (red cross) of PAM, the amidated peptide precursor (proGATI-NH_2_) and its various fragments, and the cleaved/amidated subtilisin-like endoprotease VLE1 in cilia of *minus* and *plus* vegetative and gametic cells, and in ectosomes and the soluble secretome obtained from mating gametes. PAM is present in vegetative and gametic cell cilia and is released into ectosomes but not into the secretome. In contrast, proGATI-NH_2_ is undetectable in vegetative cilia and only appears following gametogenesis. The amidated C-terminal fragment is generated in *plus* gamete cilia and released into ectosomes and the secretome during mating; other pro-GATI products are also variably present in these samples. VLE1 is found in vegetative and *plus* gamete cilia, but not in *minus* gamete cilia. All ciliary VLE1 is proteolytically processed within the pro-domain and amidated. As VLE1 moves to ectosomes and is released into the soluble secretome, it undergoes a change in domain architecture with the catalytic S8 and C-terminal domains dissociating from the amidated N-terminal segment.

With the endoproteolytic cleavage of proGATI limited to the surface of mating ectosomes (Fig. 5), we considered the possibility that its structural domains might play a role in its localization. Despite sharing little sequence similarity, a DALI search (Holm, 2020) revealed structural relatives for each proGATI domain: domain 1 is distantly similar to halohydrin dehalogenase from *Ilumobacter coccineus* (z score = 6.2, RMSD = 13.6 Å; 6I9W); domain 2 is related to EPR3 (a carbohydrate receptor) from *Lotus japonicus* (z score = 4.2, RMSD = 2.4 Å; 6QUP); and domain 3 resembles a chitosanase from *Paenibacillus sp*. (z score = 9.3, RMSD = 2.5 Å; 4ZXE). The ability of EPR3 and chitosanase to interact with carbohydrates (Kawaharada et al., 2015; Lopez-Moya et al., 2019; Wong et al., 2020) suggests that domains 2 and 3 might play a role in the tripartite interaction of proGATI with the ectosomal surface, with subsequent endoproteolytic cleavages facilitating release of specific fragments.

For signaling peptides released on ectosomes, protease resistance may be especially important. Domain 3 corresponds precisely to the 75-kDa C-ter fragment (Fig. 3E). N-glycosylation of the two potential sites in domain 3, along with O-glycosylation of a hydroxy-Pro located nine residues from the amidated C-terminus likely accounts for the ∼50 kDa discrepancy between its apparent molecular mass and the mass of its polypeptide chain (23 kDa) (Figs 3 and 8A). The endoproteolytic cleavage that produces 75-kDa C-ter fragment in ectosomes would also produce the 120-kDa N-ter fragment. Although the C-terminus of proGATI is accessible to PAM, converting its C-terminus from -GATI-Gly to -GATI-NH_2_, the amidated C-terminus is trypsin resistant (Fig. 2) and stable when exposed on the ciliary membrane and the surface of mating ectosomes.

Examination of the first *C. reinhardtii* protein known to serve as a peptide precursor indicates that it shares many similarities with vertebrate peptide precursors. However, its larger size, more complex domain organization and extensive modifications suggest that this precursor carries additional information needed to ensure that its signaling task can be accomplished.

### Controlling the endoproteolytic cleavage of proGATI

ProGATI cleavage is linked to both mating type and subcellular location (Fig. 8B). The cell bodies of *plus* and *minus* gametes contain intact proGATI, but the 75-kDa C-ter fragment is found only in the cilia of *plus* gametes. Both N- and C-ter proGATI fragments accumulate in mating ectosomes. In metazoans, the cell type-specific cleavage of propeptides such as proopiomelanocortin (Kumar et al., 2016b) and proglucagon (Drucker, 2018) reflects the cell type-specific expression of subtilisin-like prohormone convertases. Mating ectosomes contain only two subtilisin-like proteases, VLE1 and SUB14 (Luxmi et al., 2018). The presence of VLE1, but not SUB14, in the cilia of *plus* gametes, where proGATI cleavage occurs, suggests that VLE1 serves as a proGATI convertase. VLE1 is also localized to the ciliary membrane in vegetative cells (Kubo et al., 2009; Wood et al., 2013); its release into vegetative ectosomes provides access to the mother cell wall, which it degrades, allowing release of mitotic progeny. A matrix metalloproteinase (gametolysin), not VLE1, cleaves the gametic cell wall prior to fusion (Kinoshita et al., 1992), suggesting that VLE1 has additional targets on *plus* gamete cilia and/or in the extracellular milieu. VLE1 cleaves to the C-terminal side of basic residues, although the required sequence context is poorly understood (Matsuda et al., 1995). Endoproteolytic cleavage of proGATI after a basic residue within a furin-like cleavage site (R^693^FSR↓) produces the amidated 75 kDa C-ter fragment (Figs. 1F and 8A).

In metazoans, luminal pH plays a central role in controlling prohormone convertase activation and the storage of product peptides in secretory granules (Halban, 1991). With proGATI cleavage products accumulating on the surface of mating ectosomes, luminal pH cannot serve as a regulatory factor. The pro-domains of subtilisin-like endoproteases facilitate catalytic domain folding and inhibit activity. Protease activation requires autoproteolytic cleavage, separating the pro-domain from the catalytic domain (Shakya and Lindberg, 2020). Additional cleavages within the pro-domain may also be required for pro-domain release and S8 domain activation. Consistent with this, active VLE1 purified from culture medium following hatching lacked its pro-domain (Kubo et al., 2009). Our analysis of the soluble mating secretome identified the intact VLE1 S8/C-terminal domain, but not the N-terminal/pro-domain (Fig. 8B).

Sequence analysis revealed an unusual pro-domain in VLE1, with homologous sequences found only in other members of the volvocine algae (*e*.*g*. the protease VheA, required for release of juvenile *Volvox* from the parental spheroid (Fukada et al., 2006)). To understand how VLE1 activation might occur, a structural model was built using RoseTTAFold (Baek et al., 2021) (Figs. 7C and S6). The active site contains a classic Ser-His-Asp catalytic triad and an Asn residue that stabilizes the transition state in the oxyanion hole (Fig. 7C) (Shakya and Lindberg, 2020). The VLE1 pro-domain consists of an α/β fold that makes extensive contact with one face of the S8 domain. Emanating from this α/β region is an extended strand that arches over the active site, occluding it; the –Gly-Arg-Arg cleavage/amidation site is exposed on the surface. Given the large surface area buried by the pro-domain, cleavage at the –Gly-Arg-Arg site seems unlikely to result in pro-domain release from the catalytic core.

For amidation to occur, the extended strand must swing away from the catalytic site, enabling carboxypeptidases to remove remaining Arg residue(s) and allowing PAM to access the exposed Gly residue. The functional consequences of amidation at this site remain to be determined. Binding of the amidated pro-domain C-terminus to a target protein might facilitate retention of the N-terminal/pro- domain of this type II membrane protein in the ciliary membrane, allowing the enzymatically active S8/C-terminal domain to enter mating ectosomes.

### Ciliary ectosomes as an ancient mode of rapid, regulated secretion

Changes in protein expression allow unicellular organisms like *C. reinhardtii* to regulate secretion of the enzymes needed to acquire specific nutrients, but this type of response requires time. In metazoans, peptides stored in secretory granules can be released within milliseconds of signal receipt. Our data indicate that ciliary ectosomes serve as an ancient mode of rapid, regulated secretion. Like the assembly of secretory granules, the assembly of ciliary ectosomes is a highly regulated process. The cilia of both vegetative cells and mating gametes release bioactive ectosomes; their compositions are unique and developmentally regulated (Wood et al., 2013; Long et al., 2016; Cao et al., 2015; Luxmi et al., 2019). Since ectosomes are formed from the ciliary membrane, proteins targeted to ectosomes must first gain access to the cilium. The transition zone plays an essential role in establishing and maintaining the unique lipid and protein composition of the ciliary membrane (Long and Huang, 2020; Nachury and Mick, 2019; Takao and Verhey, 2016).

The entry of ciliary proteins into ectosomes is also regulated. Differences in the ectosomal trafficking of PAM, VLE1 and proGATI illustrate key features of this regulatory step (Fig. 8B). CrPAM is found in mating ectosomes but not in vegetative ectosomes (Luxmi et al., 2019); cleavage of CrPAM does not occur and active enzyme does not appear in the soluble secretome. While VLE1 is found in the cilia of *plus* gametes, the presence of the N-terminal/pro-domains, along with the intact S8/C-terminal domains suggests that ciliary VLE1 is not active. VLE1 recovered from mating ectosomes and the secretome lacks the N-terminal/pro-domains, indicating that it has been activated. While proGATI is present in the cilia of *plus* and *minus* gametes, cleavage occurs only in the cilia of *plus* gametes; more extensive cleavage of proGATI is linked to its release in mating ectosomes, where both N-ter and C-ter fragments accumulate. Although metazoan secretory granules generally store mature product peptides, the cleavage of proatrial natriuretic factor by corin, a type II plasma membrane enzyme like VLE1, is tied to the exocytosis of atrial granules (Glembotski et al., 1988).

Metazoan peptide-containing secretory granules can be stored for long periods of time, with release responding rapidly to receptor-mediated secretagogue stimulation. In *C. reinhardtii*, ciliary adhesion of mating gametes causes ectosomes to appear on the ciliary surface in a process that requires receptor-mediated signaling; strikingly, activating gametes directly with dibutyryl-cAMP does not lead to ectosome release (Cao et al., 2015). The signals that control food intake in mammals require localization of the melanocortin-4 receptor to the primary cilia of hypothalamic neurons (Wang et al., 2021). The ciliary localization of free fatty acid receptor-4 and prostaglandin-E receptor-4 in α- and β-cells plays an essential role in hormone secretion (Wu et al., 2021) and mice lacking primary cilia on their β-cells exhibit impaired glucose homeostasis and develop diabetes (Hughes et al., 2020).

As observed in mammals, multiple receptors have been identified in *C. reinhardtii* cilia (Huang et al., 2004; Luxmi et al., 2019; Ranjan et al., 2019). By taking advantage of the ease with which cilia can be isolated from *C. reinhardtii*, its precisely delineated sexual reproductive cycle and the identification of a bioactive amidated peptide in mating ectosomes, it is now clear that cilia provide a means of controlling endoproteolytic processing of propeptides and the release of mature bioactive peptide products. Although, the stimulus-dependent secretion of neuropeptides from dense core vesicles stored at the presynaptic endings of axons or exported to dendrites has been well studied (Ding et al., 2019), whether bioactive peptides are released from the primary cilia of neurons and endocrine cells remains to be determined.

In summary, this study provides a mechanism through which amidated peptide products are synthesized, post-translationally modified, trafficked into cilia and released into ciliary ectosomes by a unicellular organism, *C. reinhardtii*. As both cilia and the peptidergic signaling machinery are conserved throughout eukaryotes, this study should shed light on the mechanisms through which cilia-based secretion is regulated in health and dysregulated in various ciliopathies.

## Key Resource Table

| Reagent or Resource | Source | Identifier |
| --- | --- | --- |
| **Antibodies** |  |  |
| CrPAM luminal (rabbit) | (Kumar, 2017) | CrPHM-PAL-rhod |
| CrFMG1 (mouse) | Dr. R. Bloodgood | (Bloodgood et al., 1986) |
| ProGATI N-ter (rabbit) | This study | CT237(N-ter) |
| ProGATI C-ter (rabbit) | This study | CT237(C-ter) |
| **Chemicals, Peptides, Reagent Kits** |  |  |
| Amidated peptide, GATI-NH<sub>2</sub> | Biomatik | (Luxmi et al., 2019) |
| Control peptide, GATI-OH | Biomatik | (Luxmi et al., 2019) |
| Glycine-extended peptide, GATI-Gly | Biomatik | (Luxmi et al., 2019) |
| N-ter peptide (CYELGLDIDGKPAHPAAT-NH<sub>2</sub>) | Biomatik | This study |
| **Experimental model** |  |  |
| *Chlamydomonas reinhardtii* strains | *Chlamydomonas* resource center | CC124 and CC125 |
| HEK-293 cells | American Type Culture Collection | CRL-1573 |
| **Software** |  |  |
| Image J | NIH | https://imagej.nih.gov/ij/ |
| UNICORN FPLC software | GE Healthcare | ver. 5.2 |
| Prism 5 | GraphPad | https://www.graphpad.com |
| PyMOL | Schrödinger LLC | ver. 2.4.0 https://pymol.org |
| RoseTTAFold | Baker laboratory | https://robetta.bakerlab.org/ |
| DALI | Holm Group | https://ekhidna2.biocenter.helsinki.fi/dali/ |

## Methods

### *Chlamydomonas* cell culture and gametogenesis induction

Wild type *C. reinhardtii* mating type *minus* (CC124) and *plus* (CC125) strains were cultured in R-medium (Harris, 2009) aerated with 95% air and 5% CO_2_ under a 12 h light/12 h dark cycle at 22 °C. The strains were obtained from the *Chlamydomonas* Resource Center (https://www.chlamycollection.org/). To induce gametogenesis, vegetative cells of both mating types were washed, and resuspended in nitrogen-deficient minimal medium (M-N medium) for 24-36 h under aeration and a 12 h light/12 h dark cycle.

### Preparation of ectosomes, cilia and cell lysates from mating gametes

Gametes of both mating types were resuspended in 10 ml of fresh nitrogen-free M-N medium at a density of 5×10^6^ cells/ml. An equal number of mating type *minus* and *plus* gametes were mixed for 1 h; after incubation, ectosomes were isolated by differential centrifugation as described previously (Luxmi et al., 2019). Ectosome-enriched pellets were resuspended in TMT buffer [20 mM 2-[tris(hydroxymethyl)-methylamino]-ethanesulfonic acid (TES), pH 7.4, 10 mM mannitol, 1% Triton X-100] containing a protease inhibitor cocktail (cOmplete ULTRA Tablets, # 05892791001, Roche, Basel, Switzerland) and 0.3 mg/ml phenylmethylsulfonyl fluoride (PMSF, Sigma Chemical Co., St. Louis, MO). For electron microscopy (see below), ectosome-rich pellets were resuspended in 10 mM HEPES buffer containing the same protease inhibitors.

Cell lysates were prepared as described previously (Luxmi et al., 2019). *Minus* and *plus* mixed gametic cells were harvested by centrifugation at 1,600 xg and resuspended in TMT buffer containing 0.2 M NaCl, the protease inhibitor cocktail and 0.3 mg/ml PMSF. Gametes were deciliated using dibucaine and cilia isolated by standard methods and resuspended in HMS buffer (10 mM HEPES, pH7.4, 5 mM MgSO_4_ and 4% sucrose) (King, 1995; Witman, 1986); the deciliated cell bodies were resuspended in TMT buffer containing 0.2M NaCl, protease inhibitor cocktail and 0.3 mg/ml PMSF. Protein content was determined using the bicinchoninic acid assay (BCA) (Thermo Fisher Scientific, Rockford, IL, USA).

Samples for electrophoresis were prepared by mixing with 2× Laemmli sample buffer (Bio-rad, Hercules, California) and denatured at 55°C for 5 min; samples were fractionated in Criterion TGX 4–15% SDS-PAGE gradient gels (Bio-rad) and transferred to PVDF membranes. Proteins were visualized using Coomassie brilliant blue, the blots destained and then blocked using 5% milk dissolved in 1% Tween-20 in Tris-buffered saline. Incubation with primary antibodies was carried out overnight at 4°C; after washing, horseradish peroxidase-tagged second antibody (Thermo Fisher Scientific) was applied for 1 h at room temperature and the signal visualized using SuperSignal enhanced chemiluminescent (ECL) reagent (Thermo Fisher Scientific, #34080).

### Ciliary fractionation

Isolated cilia were incubated with TMT buffer for 60 min at 4°C, to solubilize ciliary membrane and matrix proteins. The remaining axonemes were incubated with 0.6 M NaCl in TM buffer to release axonemal proteins tightly bound *via* ionic interactions. The extracted axonemal pellet was dissolved in SDS lysis buffer (0.5% (w/v) sodium dodecyl sulfate, 0.05 M Tris.Cl, pH 8.0) containing protease inhibitor cocktail and 0.3 mg/ml PMSF. Soluble samples were desalted and concentrated using Amicon concentrators (10-kDa cut-off; Millipore Sigma, # UFC800308; Merck KGaA, Darmstadt, Germany). Samples (20 µg protein) were fractionated by SDS-PAGE and analyzed by immunoblotting.

### Immunofluorescence microscopy

Resting gametes were harvested by centrifugation at 1,600 ×g and fixed with 2% paraformaldehyde in buffer containing 30 mM HEPES, 5 mM EGTA, 5 mM MgSO_4_, 25 mM KCl, 4% sucrose, pH 7.0. Cells were allowed to adhere to 0.1% polyethyleneimine-coated coverslips for 10 min and then treated with methanol for 10 min at −20°C. Subsequent blocking and antibody incubation were done as described (Luxmi et al., 2019). Primary antibodies used were affinity-purified rabbit N-ter and C-ter proGATI antibodies (from CT327; 1:500) and mouse FMG1 (1:1000). Alexa 488 anti-rabbit (Life Technologies, Thermo Fisher Scientific) (1:500) and Cy3 anti-mouse (Jackson ImmunoResearch Laboratories, West Grove, PA) (1:2000) conjugates were used as secondary antibodies. Images were obtained using a Zeiss 880 confocal microscope with a 63× oil objective.

### Electron microscopy analysis

Immuno-gold labeling of ectosomes was performed as described previously (Luxmi et al., 2019) with the following modifications. Freshly isolated mating ectosomes were fixed with 1% paraformaldehyde (EM grade) and incubated on ice for 30 min. Fixed samples were placed on glow-discharged 400-mesh carbon-coated nickel grids (Electron Microscopy Sciences, Hatfield, PA) for 10-20 min and then washed with 1x PBS (137 mM NaCl, 10 mM phosphate, 2.7 mM KCl pH 7.4) and incubated with 50 mM glycine. Samples were incubated overnight at 4°C with affinity-purified N-ter and C-ter antibodies (1:10), washed and incubated for 1 h at room temperature with gold conjugated (10-nm) goat anti-rabbit-IgG (1:15, Electron Microscopy Sciences).

For thin section EM, freshly isolated ectosomes were fixed with 2.5% glutaraldehyde in 0.1 M cacodylate buffer, pH 7.4 for 1 h at 4°C. After fixation, ectosomes were centrifuged at 424,000 xg for 30 min and the ectosome pellet was washed with 0.1 M cacodylate buffer, pH 7.4. Pellets were then transferred to 0.5 ml tubes; after the buffer was carefully removed, ultra-low gelling agarose (100 μl of 4%) was added and the sample was immediately centrifuged at 1,600 xg for 10 min at room temperature. Tubes were then placed on ice for 10 min to solidify the agarose. Ultra-thin sections of agarose-embedded ectosomes were mounted on 200-mesh copper/rhodium grids, and imaged using a H-7650 transmission EM (Hitachi High Technologies Corporation, Tokyo, Japan) operating at 80 kV.

### PreproGATI in HEK-293 cells

HEK-293 cells were maintained in DMEM/F12 medium containing 10% fetal calf serum (Hyclone), 100 units/ml penicillin-streptomycin and 25 mM HEPES, pH 7.4 at 37°C in a 5% CO_2_ incubator. A cDNA (2742 bp) encoding preproGATI was synthesized and cloned into pUC57 (GenScript). This cDNA was then subcloned into pCI-neo (Promega, Madison, WI) and verified by sequencing. Transient transfections were performed using lipofectamine 3000 (Invitrogen, Thermo Fisher Scientific) and stable populations of HEK-293 cells expressing preproGATI were generated by selecting cells in DMEM/F12 medium containing 0.5 mg/ml G418 disulfate (KSE Scientific, Durham, NC). For analyzing spent medium, cells were washed with serum-free medium [DMEM/F-12 medium containing insulin-transferrin-selenium (ITS) (Thermo Fisher Scientific), 25 mM HEPES, pH 7.4, 100 units/ml penicillin-streptomycin, 1 mg/ml BSA] and then incubated in serum-free medium at 37°C with 5% CO_2_. Cell lysates were prepared in 1x SDS lysis buffer with 1x protease inhibitor cocktail (Sigma, # P8340) and 0.3 mg/ml PMSF. Soluble fractions (equal protein) were analyzed using standard electrophoretic and immunoblotting techniques.

### HEK-proGATI purification

Stably transfected HEK-293 cells expressing preproGATI (HEK-GATI cells) were washed and cultured in serum-free media lacking ITS and BSA for 16-18 h. Spent medium was collected and centrifuged at 100 xg to remove cell debris. Protease inhibitor cocktail and 0.3mg/ml PMSF were added to the medium, which was stored at −80°C. Spent medium pooled from multiple sequential collections was used for purification. A weak anion exchange column, HiTrap ANX Sepharose FF (Cytiva # 17-5163-01; Sigma), was used to concentrate the HEK-proGATI (pI = 6.04). Prior to sample loading, the pH of the spent medium was adjusted to 7.5 and the sample centrifuged at 10,000 xg for 15 min to remove any insoluble material. The HiTrap ANX Sepharose FF column was washed with water, and equilibrated with 20 mM Tris, pH 7.5 containing 100 mM NaCl and 5% glycerol. The sample was loaded with a peristaltic pump and the flow-through discarded. The column was washed with 20 mM Tris, pH 7.5 buffer containing 100 mM NaCl and 5% Glycerol until the phenol red from the spent medium was no longer visible. Proteins were then eluted using an AKTA Purifier 10 FPLC System (GE Healthcare, Fairfield, CT), with a gradient of 100 mM to 1 M NaCl in 20mM Tris buffer containing 5% glycerol, a flow rate of 1 ml/min and a total elution volume of 40 ml. The collected fractions were analyzed using 4-15% SDS-PAGE gels, immunoblotted and probed with the C-ter antibody. Peak fractions were pooled and further purified by gel filtration using a Superdex 200 Increase 10/300 GL (GE Healthcare, 28-9909-44) column equilibrated with 20 mM HEPES, pH 7.4 containing 0.5 M NaCl (Fig. S2C). Fractions were pooled based on SDS-PAGE analysis; purified HEK-proGATI (∼5 µg) was then analyzed by mass spectrometry (see below). Approximately 5 mg of HEK-proGATI was purified from 500 ml of spent medium.

### BCS treatment of HEK-GATI cells

HEK-GATI cells plated into 24 well dishes were washed and incubated for 30 min in serum-free media, at 37°C with 5% CO_2_. Cells were then treated with serum-free media containing 50 µM bathocuproine disulfonic acid (BCS, Sigma) as described by (Bonnemaison et al., 2015). Cells treated with medium only were used as control. Spent medium was collected and centrifuged at 100 xg to remove cell debris. Cell lysates (15 μg, ∼20% of total) and spent media (15 µl, 5% of total) were fractionated in 4-15% SDS-PAGE gels and analyzed by immunoblotting.

### Antibody generation

Synthetic peptides (BioMatik, Kitchener, Ontario, Canada) from the N-terminal (YELGLDIDGKPAHPAAT-NH_2,_ 1.5 mg) and C-terminal (YAPGTGGGATI-NH_2_, 1.5 mg) regions of proGATI were individually conjugated to keyhole limpet hemocyanin (KLH; 3 mg; Sigma H-7017, Lot 110K4833). An additional Cys residue added to the N-ter peptide allowed conjugation to KLH using m-maleimidobenzoyl-N-hydroxysuccinimide ester. KLH conjugation of the C-ter peptide used glutaraldehyde, facilitating the generation of amide specific antibodies. Three rabbits (CT327, CT330, and CT332) were immunized with a mixture of KLH-conjugated N-ter and C-ter peptides by Covance Immunology Services (Denver, PA). Crude IgG was obtained by ammonium sulfate precipitation from the sera of immunized rabbits and N-ter and C-ter antibodies further purified by peptide affinity chromatography. The N-ter (pI 5.5) and C-ter (pI 9.9) peptides were conjugated to Affi-Gel-10 (Bio-rad) agarose beads for affinity-purification. Recoveries during affinity purification and cross-reactivity of purified antibodies were examined using solid phase assays. High-affinity binding 96-well plates coated with N-ter (5 ng) or C-ter (5 ng) peptide were prepared and serial 3-fold dilutions of each sample were tested.

### Deglycosylation assays

The presence of N-linked oligosaccharides was examined using PNGase F (New England Biolabs (NEB), Ipswich, MA, # P0708S) and the presence of O-linked sugars was assessed by combined treatment with O-glycosidase (NEB #P0733S) and α2-3,6,8 neuraminidase (NEB #P0720S). Mating *C. reinhardtii* ectosomes (20 µg) and spent medium (9 µl) from HEK-293 cells expressing proGATI were denatured by heating at 100°C for 10 min with 1x denaturing buffer. Following denaturation, samples were deglycosylated following the manufacturer’s protocol. Samples incubated on ice only (-) and treated with buffer only (+B) were used as controls. For mass spectrometry analysis (see below), purified HEK-proGATI (∼5 μg) was denatured and deglycosylated using deglycosylation mix II (NEB, #P6044S), which contains the enzymes needed to remove N-linked and many common O-linked glycans. The deglycosylated sample was buffer exchanged using Zeba™ spin desalting columns (40K Mol. Wt. cutoff; Thermo Fisher Scientific, #87768).

### Immunoprecipitation

Immunoprecipitation was performed using with slight modifications of previous protocols (Miller et al., 2017). Cross-reactive proteins were immunoprecipitated from *plus* gametic cell lysates and from mating ectosomes using affinity-purified C-ter antibody. Before immunoprecipitation, samples were denatured. An equal volume of 1× SDS-P buffer (50 mM Tris pH 7.6, 1% SDS, 130 mM NaCl, 5 mM EDTA, 50 mM NaF, 10 mM NaPP_i_) containing 0.3 mg/ml PMSF, protease inhibitor cocktail and PhosStop (Roche) was added to the TMT cell lysate (1 mg protein) or mating ectosomes (1 mg protein) and samples were heated at 55°C for 5 min. Samples were allowed to cool and incubated with 0.5 volume (for cell lysate) or 1.0 volume (for ectosomes) of 15% NP-40 for 20 min on ice. Samples were then diluted with 5 volumes of TES-mannitol (TM) buffer containing protease inhibitor cocktail, 0.3 mg/ml PMSF and PhosStop. Each sample was centrifuged at 15,000 xg for 15 min at 4°C to remove any insoluble material. For pre-clearing, washed Protein A agarose beads (50 µl) (Thermo Fisher Scientific, #22810) were added, samples were tumbled for 30 min at 4°C and then centrifuged at 100 xg for 3 min. Affinity-purified C-terminal antibody (100 µl) was then added to the pre-cleared supernatants, followed by Protein A beads (50 μl) that had been washed with 1x TMT buffer containing 1x protease inhibitor cocktail, 0.3 mg/ml PMSF and 1x Phos Stop. After overnight incubation at 4°C, beads were pelleted and the unbound fraction saved; beads were then washed once with TMT buffer containing 0.5M NaCl and twice with TM buffer containing protease inhibitor cocktail, 0.3 mg/ml PMSF and Phos Stop. Bound protein was eluted by boiling in 2x Laemmli sample buffer (Bio-rad) and particulate material removed by centrifugation at 15,000 xg at room temperature. The input (15 µg) and eluted proteins (∼2% of IPT) were fractionated in 4−15 % SDS-PAGE gels (Bio-rad) and analyzed by immunoblotting. For mass spectrometry, samples were fractionated by SDS-PAGE and visualized using QC colloidal Coomassie stain (Bio-rad); the 75-kDa band was excised from the *plus* gamete cell lysate and mating ectosome samples.

### Mass spectrometry

Excised gel bands were destained using 40% ethanol and 10% acetic acid in water, equilibrated to pH 8 in 100 mM ammonium bicarbonate, reduced by incubation with 10 mM dithiothreitol in 100 mM ammonium bicarbonate (1 hr at 37°C) and alkylated by incubation with 55 mM iodoacetamide in 100 mM ammonium bicarbonate (45 min at 37°C in the dark). Gel bands were dehydrated using acetonitrile, dried, and then rehydrated in a 12.5 ng/µL trypsin solution (Promega porcine sequencing grade trypsin) in 100 mM ammonium bicarbonate. Proteolysis proceeded for 16 hr at 37°C. Tryptic peptides were extracted using alternating washes with 100 mM ammonium bicarbonate and 5% formic acid in 50% acetonitrile and a final wash cycle with 100 mM ammonium bicarbonate and 100% acetonitrile. Peptide solutions were pooled, dried and peptides resuspended in 0.1% formic acid in water prior to mass spectrometry analysis.

Purified HEK-proGATI was diluted with 100 mM ammonium bicarbonate in water and subjected to reduction and alkylation using 5 mM dithiothreitol in 100 mM ammonium bicarbonate (1.5 hr at 37°C) and 10 mM iodoacetamide in 100 mM ammonium bicarbonate (45 min at 37°C in the dark), respectively. Promega sequencing grade trypsin was added (1:20 w/w, enzyme:protein) and proteolysis proceeded for 16 hr at 37°C. Digestion was quenched by addition of concentrated formic acid. Peptides were desalted using high capacity C_18_ desalting spin columns (Pierce #89851; ThermoFisher). Desalted peptides were dried to completion and resuspended in 0.1% formic acid in water prior to mass spectrometry analysis.

Resuspended peptides were analyzed using nanoflow ultra-high performance liquid chromatography (UPLC) coupled to tandem mass spectrometry (MS/MS) using a Dionex Ultimate 3000 RSLCnano UPLC system and Q Exactive HF mass spectrometer (ThermoFisher Scientific). Peptides were loaded onto a 75 µm x 25 cm nanoEase m/z Peptide BEH C_18_ analytical column (Waters Corporation, Milford, MA), separated using either a 1 or 2 hr reversed-phase UPLC gradient, and directly ionized into the Q Exactive HF using positive mode electrospray ionization. MS/MS data were acquired using a data-dependent Top15 acquisition method. All raw data were searched against the *C. reinhardtii* proteome using the following variable modifications: <u>Modification set 1</u> - Met and Pro oxidation, Ser, Thr, and Tyr phosphorylation, Glu, Asp, peptide C-term amidation, Cys carbamidomethylation, and Asn, Ser, Thr HexNAcylation, or <u>Modification set 2</u> - Met and Pro oxidation, Glu, Asp and peptide C-term amidation, Cys carbamidomethylation, and the following on Pro residues: 1Hyp1Hex0Pent, 1Hyp2Hex0Pent, 1Hyp3Hex0Pent, 1Hyp4Hex0Pent, 1Hyp0Hex1Pent, 1Hyp0Hex2Pent, 1Hyp0Hex3Pent, 1Hyp0Hex4Pent, 1Hyp1Hex1Pent, 1Hyp1Hex2Pent, 1Hyp1Hex3Pent, 1Hyp1Hex3Pent, 1Hyp2Hex0Pent, 1Hyp2Hex1Pent, 1Hyp2Hex2Pent, 1Hyp3Hex1Pent where Hyp = Hydroxyproline, Hex = hexose, Pent = pentose. Trypsin C-terminal cleavage specificity was set to “semi-specific C-ragged” at “KR” sites to identify C-terminal non-tryptic proteolysis sites and subsequent C-terminal peptide amidation. Peptide output option was set to “automatic score cut” to allow 0-5% peptide level FDR filtering and protein FDR was set to 1%. All other parameters were kept at default settings. Scaffold v4 or v5 (Proteome Software, Inc., Portland, OR) were used for visualization and further analysis.

For comparative proteomics of VLE1 and SUB14, vegetative and gametic cilia were obtained from both mating types by the dibucaine method (see above). Isolated cilia were separated into membrane/matrix and axonemal fractions by extraction with 1% IGEPAL CA-630 and differential centrifugation. Samples were electrophoresed in triplicate using a short SDS-PAGE gel protocol, stained with Coomassie blue and then subject to tryptic digestion. Tandem MS/MS spectra of purified tryptic peptides were obtained at the University of Massachusetts Medical School mass spectrometry facility and analyzed using Mascot with a parent ion tolerance of 10.0 ppm and a fragment tolerance of 0.050 Da. Modifications allowed included carbamidomethyl on Cys, C-terminal minus Gly plus amide, N-terminal pyroglutamylation, methionine oxidation, N-terminal acetylation and phosphorylation.

### Bioinformatics analysis and structural modeling

The signal peptide was identified using Signal P (www.cbs.dtu.dk/services/SignalP/) and N-glycosylation sites were predicted with NetNGlyc (www.cbs.dtu.dk/services/NetNGlyc/). The structural models for proGATI and VLE1 were generated using RoseTTAFold (https://robetta.bakerlab.org) (Baek et al., 2021). Structures were displayed using PyMOL (Schrödinger LLC). Structural homologues of individual proGATI domains were identified using DALI (http://ekhidna2.biocenter.helsinki.fi/dali/; (Holm, 2020)).

### Statistics and quantification

For each experiment, the number of biological replicates is indicated in the Figure Legend. One-way ANOVAs with Tukey’s multiple comparison test and two-way ANOVAs with Bonferroni post-tests were used to compare the means. Results are represented as mean ± SEM or ± range as indicated in the Figure Legend. GraphPad Prism 5 software was used to perform all statistical analyses.

## Acknowledgements

We gratefully acknowledge the quantitative proteomics analysis conducted by Dr. Jeremy L. Balsbaugh, Director of the Proteomics & Metabolomics Facility, a component of the Center for Open Research Resources and Equipment at the University of Connecticut. We also thank Maya Yankova for assistance with electron microscopy, Dr. Miho Sakato-Antoku for preparing vegetative and gametic cilia samples for mass spectrometry and assistance with chromatography, and Dr. R. Bloodgood (University of Virginia) for the gift of FMG-1 antibody.

## Supplemental Figures

**Figure S1.**
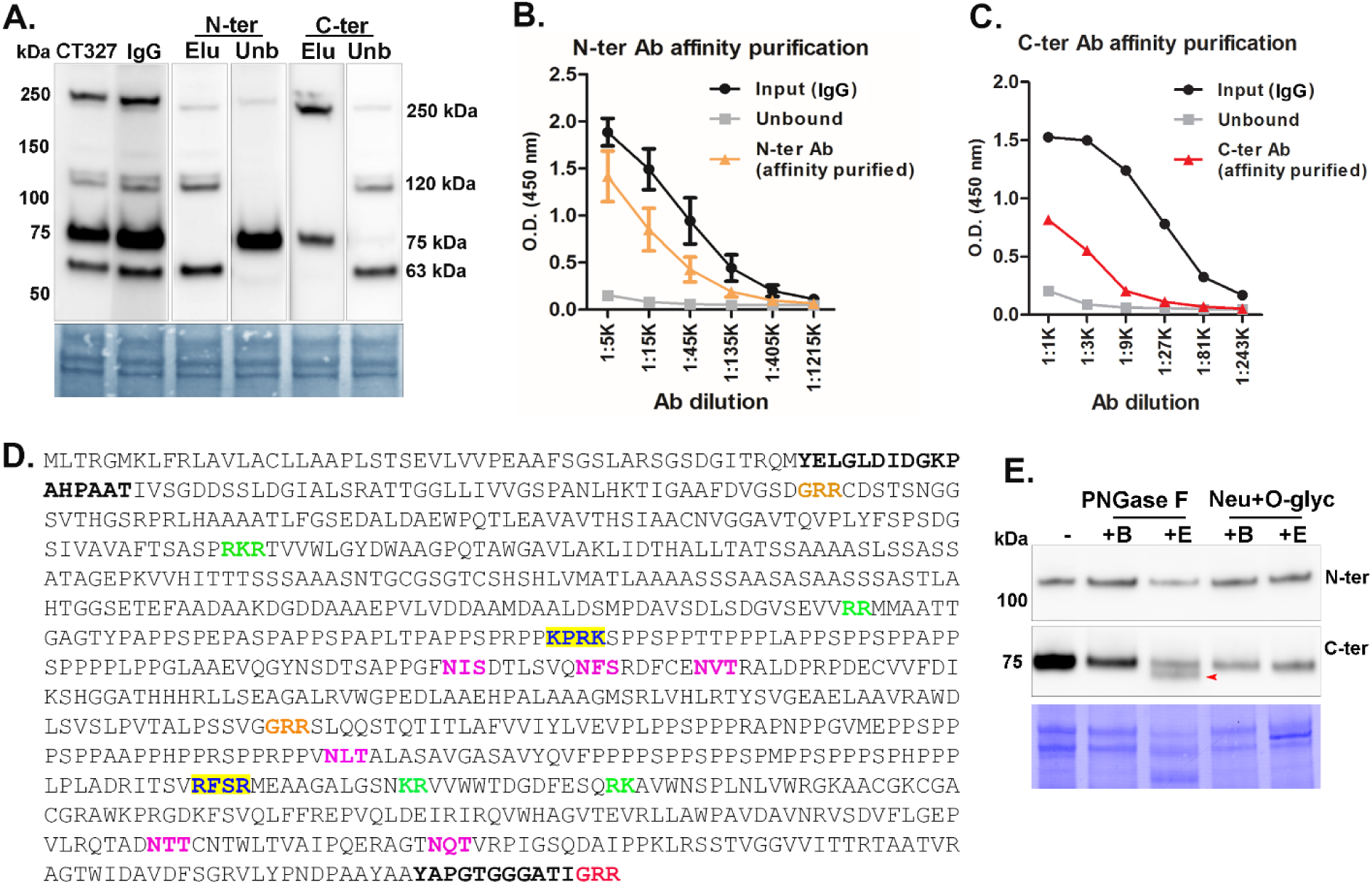
**A**. Immunoblots showing the results of affinity-purification of N-ter and C-ter proGATI antibodies. Mating ectosomes (15 µg protein) were fractionated by SDS-PAGE and individual PVDF strips incubated with serum from rabbit CT327, an immunoglobulin-enriched fraction prepared from this serum (IgG) or aliquots of the material that did not bind (unbound (Unb)) or was eluted from (Elu) columns that contained either the N-ter or C-ter peptide linked to AffiGel. The 250-kDa, 120-kDa and 63-kDa bands were detected by the bound (affinity-purified) fraction of the N-ter antibody and by the unbound fraction of the C-ter antibody. The 250-kDa and 75-kDa bands were detected by the bound (affinity-purified) fraction of the C-ter antibody and by the unbound fraction of the N-ter antibody. **B** and **C**. The specificity and yield of the affinity-purified N-ter and C-ter antibodies was determined using a solid phase assay (ELISA) with the N-ter or C-ter peptide, respectively, bound to the plate. The IgG-enriched input (IgG), unbound, and affinity-purified N-ter and C-ter antibodies were tested. **D**. The sequence of preproGATI is shown. The N-ter and C-ter antigenic peptides are indicated in bold (black); the identified C-ter amidation site is in red and other potential cleavage/amidation sites are in orange; paired basic cleavage sites (green), furin-like cleavage sites (blue with yellow highlight) and six potential N-glycosylation sites (-NXS/T; pink) are marked. **E**. Mating ectosomes (15 µg protein) were analyzed without treatment (-) or after digestion with PNGase F (buffer alone, +B; with enzyme, +E), revealing a reduction in molecular mass (red arrow) of the 75-kDa C-ter product but not the 120-kDa N-ter proGATI product. Digestion with neuraminidase and O-glycosidase (buffer alone, +B; with enzyme, +E) had no effect. The results were replicated in independent experiments.

**Figure S2.**
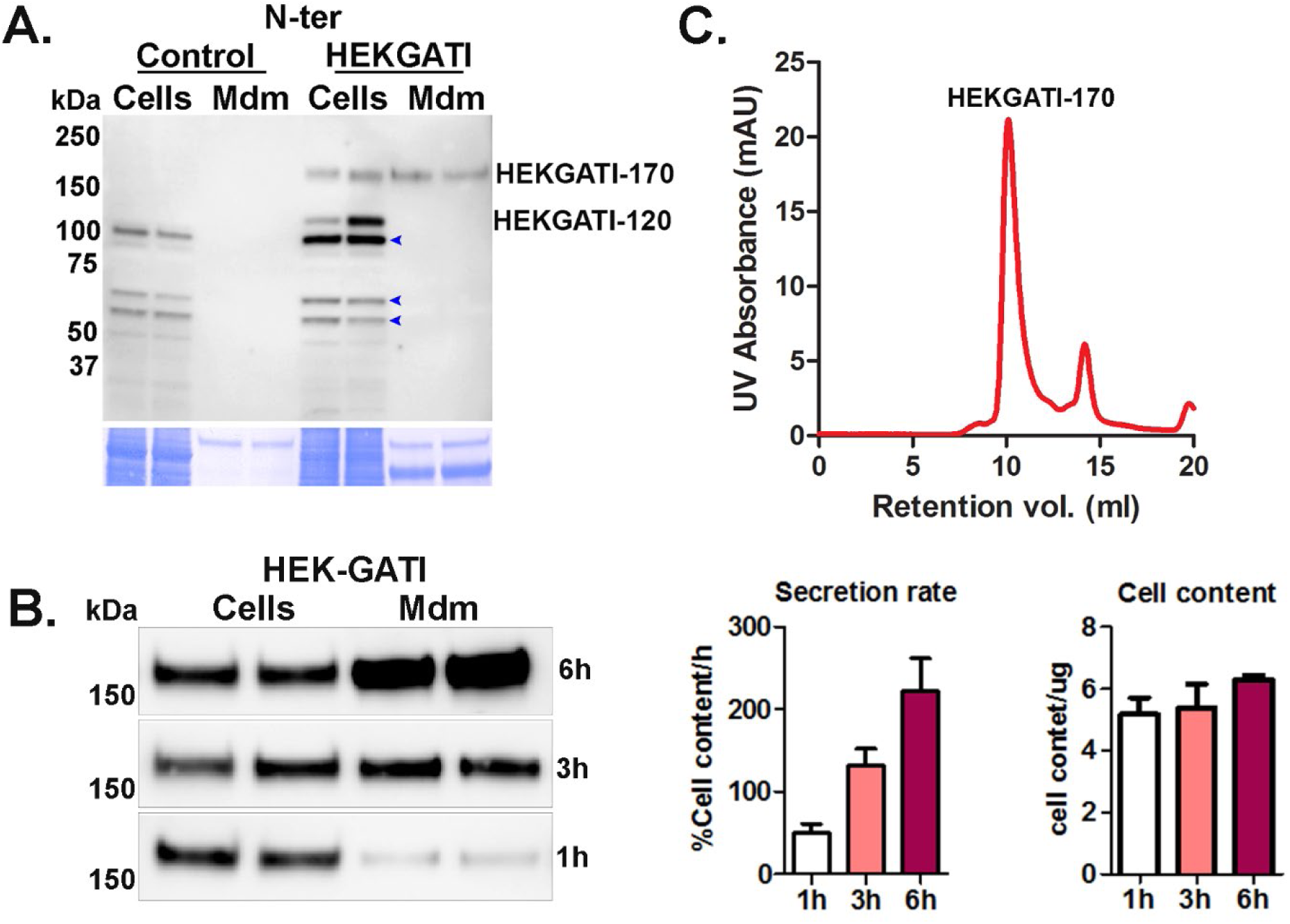
**A**. Cell extracts (Cells) and the spent medium (Mdm) from HEK-preproGATI cells and Control cells were probed with the N-ter proGATI antibody. HEK-proGATI protein (HEK-GATI-170) was detected in both cell lysates and spent media, while HEK-GATI-120 was only present in cell extracts. Non-specific bands (present in Control) are marked with blue arrowheads. Equal amounts of cell lysate (20 μg, 10% of total) and spent medium (1% of total from an 18 h collection) were analyzed. **B**. HEK-GATI cells were incubated in serum-free medium for 1, 3 and 6 h time periods; cell lysates and spent media were analyzed with the C-ter antibody. The HEK-GATI cells secrete the 170-kDa product rapidly; its secretion rate increased over a period of 6 hours (left graph), while the cell content remained constant (right graph). **C**. Gel filtration chromatogram of purified HEK-proGATI protein. The relative absorbance, measured at 280 nm (red line) shows the peak of purified HEK-proGATI protein.

**Figure S3.**
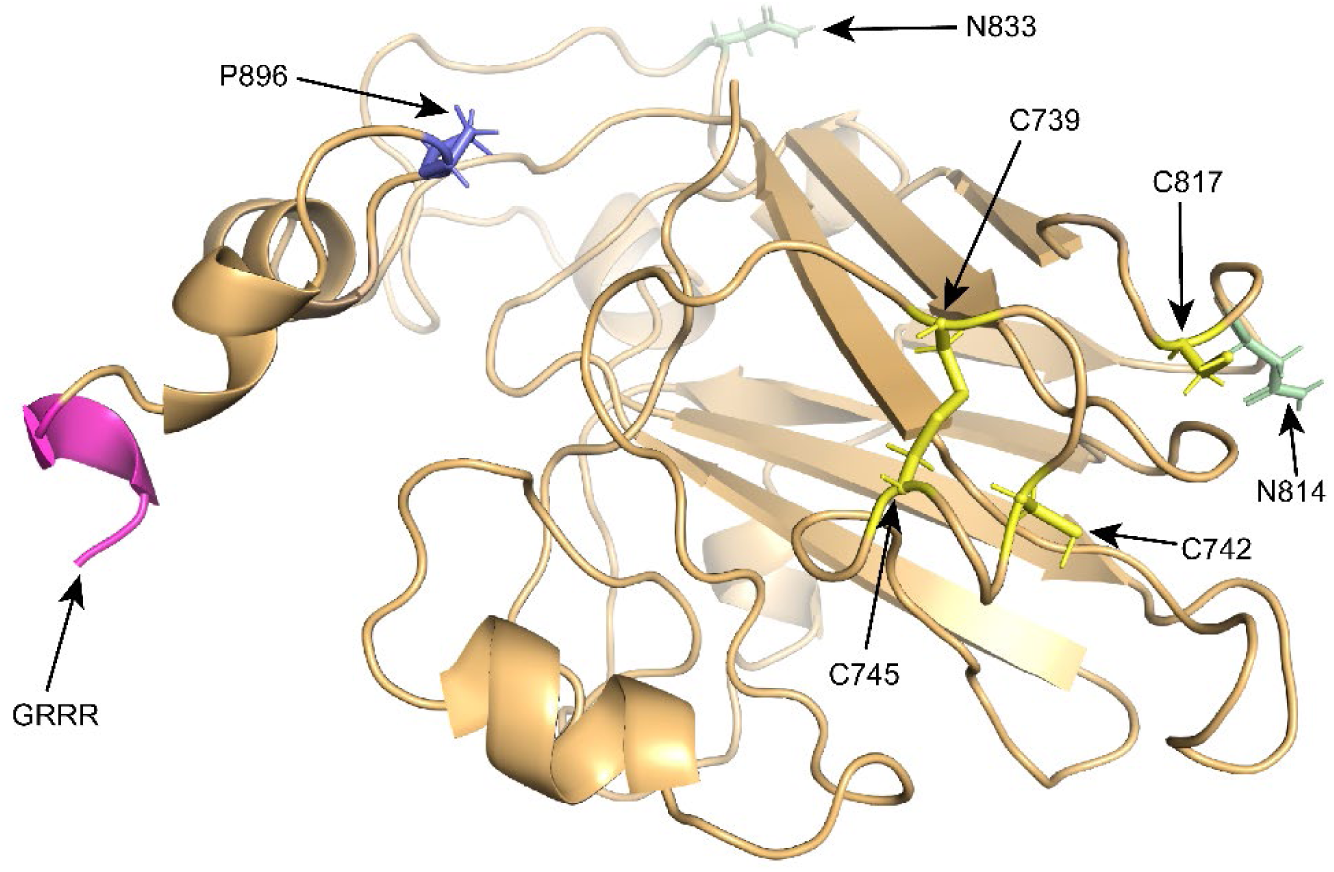
The detailed structural model for domain 3 of proGATI is shown. This domain forms an antiparallel β sandwich. The post-translationally modified residues identified in purified HEK-proGATI protein are all exposed on the surface; two deamidated Asn residues (marked in green), one HyP residue (purple) and the C-terminal amidation site (pink) are indicated. This domain contains four Cys residues (indicated in yellow); two (C^739^ and C^745^) are predicted to form a disulfide bond, while two others (C^742^ and C^817^), although in relatively close proximity, are not.

**Figure S4.**
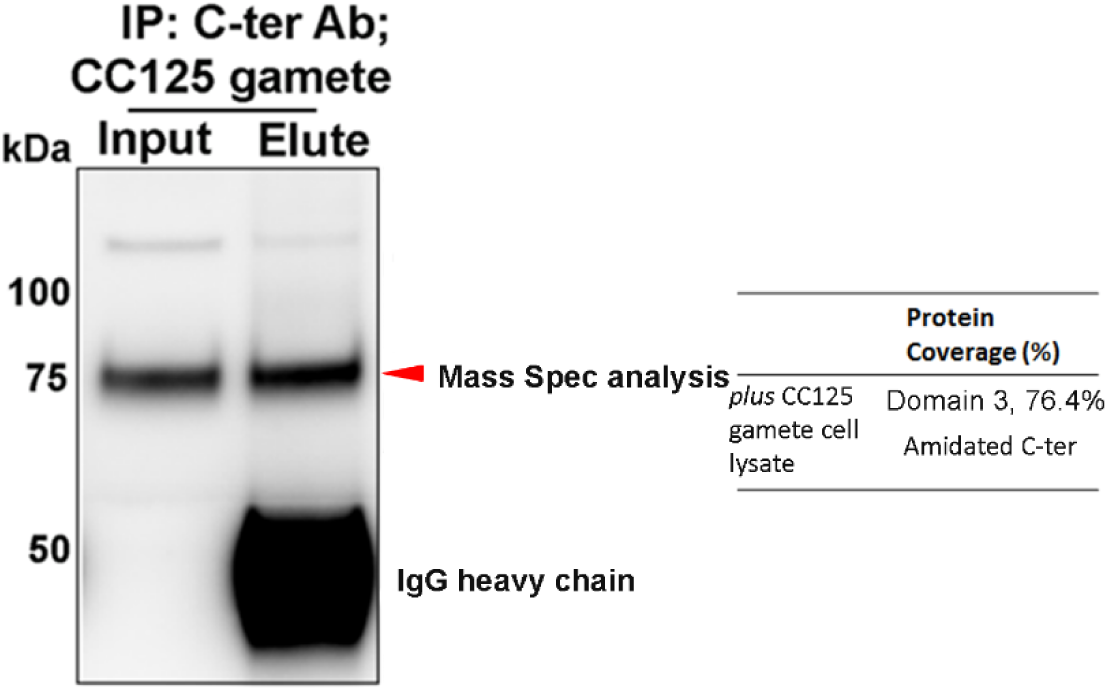
Affinity-purified C-ter antibody was used to immunoprecipitate cross-reactive material from *plus* gamete cell lysates. The 75-kDa fragment (red arrowhead) was analyzed by mass spectrometry. Only peptides from the C-terminal region of proGATI were identified; all were to the C-terminal side of the furin-like cleavage site illustrated in Fig. 1E. The C-terminal peptide (VLYPNDPAAYAAYAPGTGGGATI-NH_2_) identified in *plus* gametes was α-amidated.

**Figure S5.**
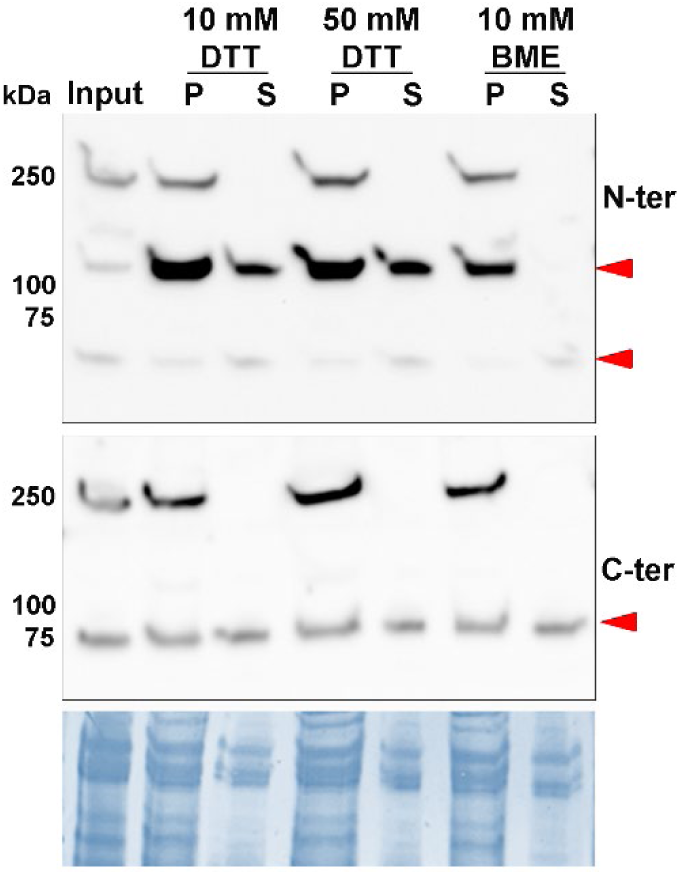
Isolated ectosomes (input) were washed with buffer containing either 10 or 50 mM dithiothreitol (DTT) or 10 mM β-mercaptoethanol (BME). Although the 120-kDa N-terminal product was released into the supernatant by DTT, it remained ectosome-associated in the presence of β-mercaptoethanol. No differential effects of these reagents were observed for the 75-kDa C-terminal fragment that is released from ectosomes by buffer treatment alone (see Fig. 6C).

**Figure S6.**
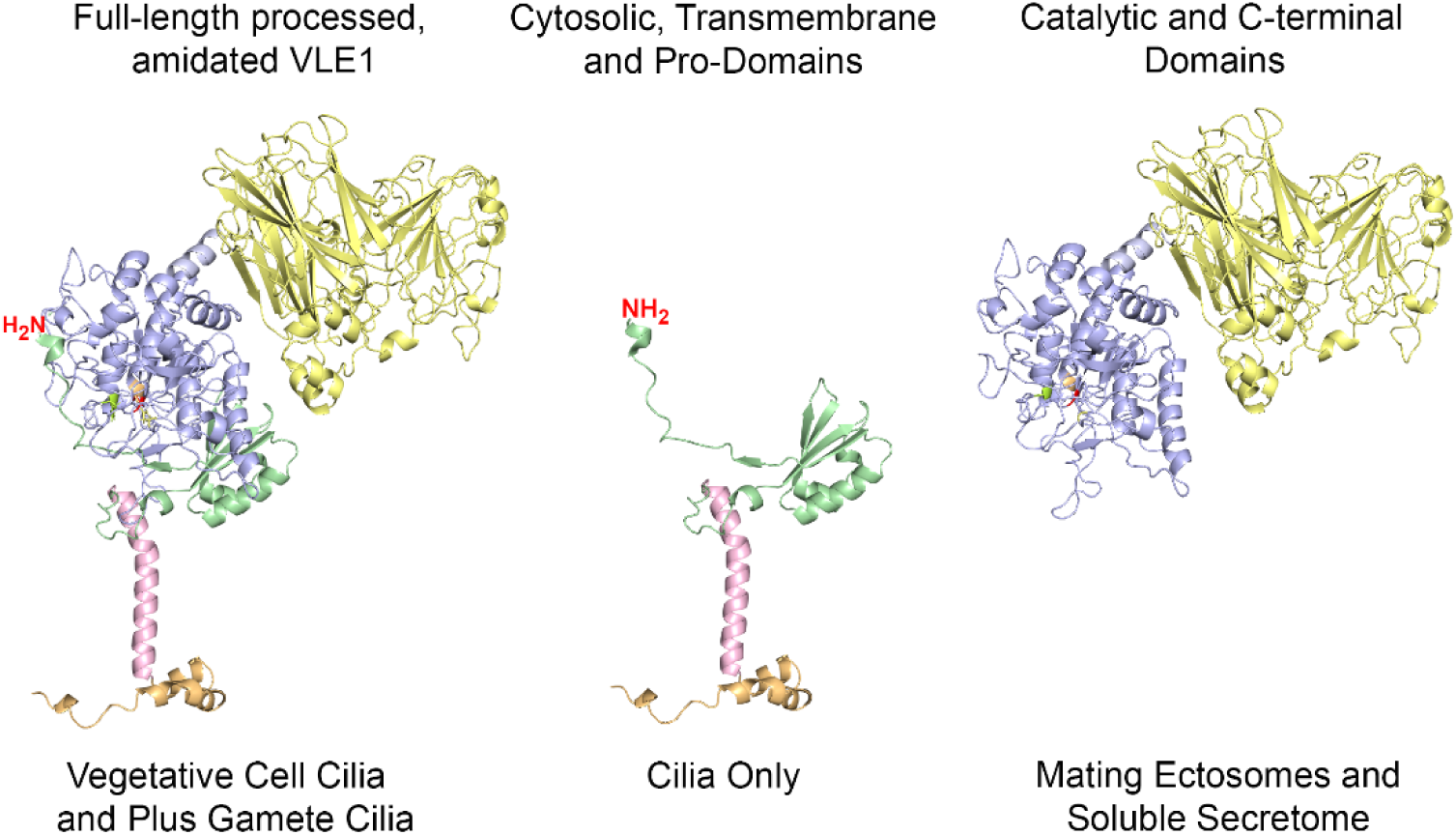
Changes in VLE1 domain organization occur during trafficking of VLE1 from cilia to ectosomes and the soluble secretome. When on the ciliary membrane of both vegetative cells and plus gametes, full-length VLE1 that has been cleaved at the diarginine site and subsequently amidated is present. However, as VLE1 moves to ectosomes and is released into the secretome, the catalytic S8 and C-terminal domains dissociate from the pro-domain which is not present in these fractions; the N-terminal/pro-domain region is presumably either retained on the ciliary membrane or trafficked back to the cell body for degradation.

## Supplemental Data

**Supplemental Data File 1.**
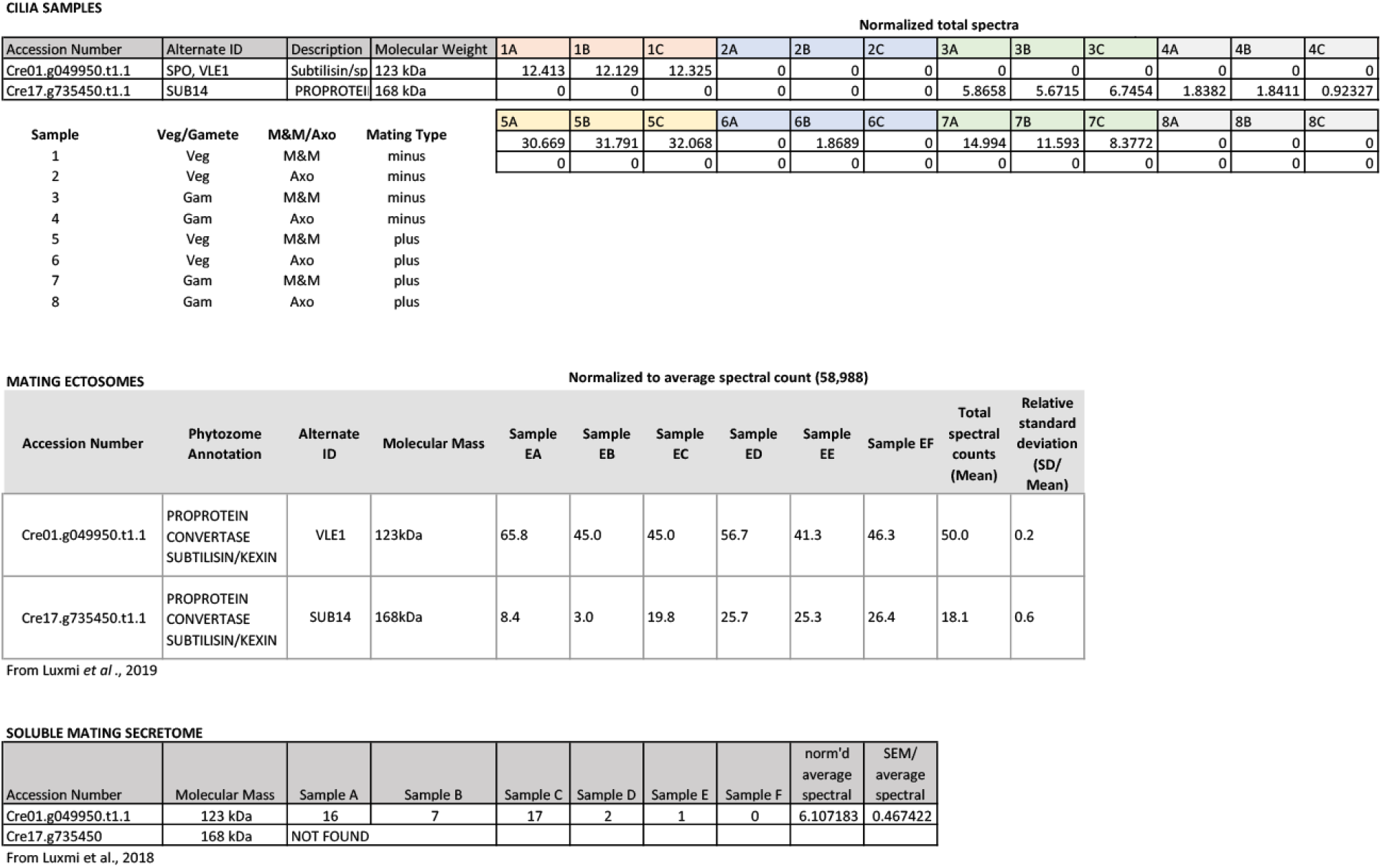
Mass spectral data (normalized total spectral counts) for VLE1 and SUB14 in vegetative and gametic cilia of both mating types. Cilia were fractionated into a detergent-soluble membrane plus matrix fraction and an axonemal fraction. All samples were analyzed in triplicate. Also shown are the normalized average spectral counts for the presence of these two proteins in mating ectosomes (data from (Luxmi et al., 2019)) and soluble mating secretome (data from (Luxmi et al., 2018)) (*.xlsx format).

### Source Data File

This file contains annotated uncropped gel and blot images used to prepare Figs. 1B, 1C, 1D, 2A, 2B, 2D, 2E, 3A, 4A, 5B, 5D, 5E, 6A, 6C, S1A, S1E, S2A, S2B, S4, and S5. (*.docx format).

